# A homologue of the mammalian tumour suppressor protein PTEN is a functional lipid phosphatase and required for chemotaxis in filamentous fungi

**DOI:** 10.1101/2020.09.15.298703

**Authors:** Berit Hassing, Alyesha Candy, Carla J. Eaton, Tania R. Fernandes, Carl H. Mesarich, Antonio Di Pietro, Barry Scott

## Abstract

Phosphoinositides (PI) are essential components of eukaryotic membranes and function in a large number of signalling processes. While lipid second messengers are well studied in mammals and yeast, their role in filamentous fungi is poorly understood. We used fluorescent PI-binding molecular probes to localise the phosphorylated phosphatidylinositol species PI[3]P, PI[3,5]P_2_, PI[4]P and PI[4,5]P_2_ in hyphae of the endophyte *Epichloë festucae* in axenic culture and during interaction with its grass host *Lolium perenne*. We also analysed the roles of the phosphatidylinositol-4-phosphate 5-kinase MssD and the predicted phosphatidylinositol-3,4,5-triphosphate 3-phosphatase TepA, a homologue of the mammalian tumour suppressor protein PTEN. Deletion of *tepA* in *E. festucae* and in the root-infecting tomato pathogen *Fusarium oxysporum* had no impact on growth in culture or the host interaction phenotype. However, this mutation did uncover the presence of PI[3,4,5]P_3_ in septa of *E. festucae* and showed that TepA is required for chemotropism in *F. oxysporum*. The identification of PI[3,4,5]P_3_ in septa of *ΔtepA* strains suggests that filamentous fungi are able to generate PI[3,4,5]P_3_ using an alternative biosynthetic pathway and that fungal PTEN homologues are functional lipid phosphatases. The *F. oxysporum* chemotropism defect demonstrates a conserved role of PTEN homologues in chemotaxis across protists, fungi and mammals.

## Introduction

Phosphoinositides (PI) regulate key cellular functions by interacting with proteins that reside in membranes or by recruiting proteins to membranes through PI-binding domains or polybasic amino acid (aa) stretches (Balla, 2013; De Craene et al., 2017). The asymmetric distribution of PIs within organelle membranes defines and maintains membrane identity (De Craene et al., 2017; Falkenburger et al., 2010). PIs consist of a glycerol backbone with two fatty acid chains attached via an ester bond, and a headgroup consisting of phosphate esterified to an inositol ring. The ring can be phosphorylated or dephosphorylated at positions 3, 4 or 5 by specific kinases and phosphatases, respectively, resulting in seven phosphorylated PI species: PI[3]P, PI[4]P, PI[5]P, PI[3,4]P_2_, PI[3,5]P_2_, PI[4,5]P_2_, PI[3,4,5]P_3_ (Balla, 2013; De Craene et al., 2017). To date, only mammals have been shown to produce all seven PIs, while in plants, neither PI[3,4]P_2_ nor PI[3,4,5]P_3_, and in fungi, neither PI[5]P nor PI[3,4]P_2_, have been detected (De Craene et al., 2017).

The most abundant PI is PI[4,5]P_2_, which accounts for 30% of the PIs in yeast and 45% in humans, and localises predominantly at the inner leaflet of the plasma membrane (PM) (De Craene et al., 2017; Di Paolo and De Camilli, 2006). PI[4,5]P_2_ plays essential roles in endo-and exocytosis, actin cytoskeleton rearrangement, the activity of ion channels and the establishment of cell polarity (Balla, 2013; De Craene et al., 2017). In mammalian cells, PI[4,5]P_2_ is generated via phosphorylation of either PI[4]P or PI[5]P by type I phosphatidylinositol 4-phosphate 5-kinases or type II phosphatidylinositol 5-phosphate 4-kinases, respectively. In addition, PI[4,5]P_2_ is also produced through dephosphorylation of PI[3,4,5]P_3_ by a phosphatidylinositol 3,4,5-trisphosphate 3-phosphatase named PTEN (Phosphatase and TENsin homologue). PTEN acts both as a lipid phosphatase and as a dual specificity tyrosine-, serine-and threonine-protein phosphatase. The lipid phosphatase activity of mammalian PTEN is essential for its role as a tumour suppressor protein, as dephosphorylation of PI[3,4,5]P_3_ inhibits the activation of downstream components of the phosphatidylinositol PI3K/AKT pathway (Manning and Cantley, 2007). As a consequence, disruption of PTEN results in deregulated cell proliferation and the onset of cancer (Chen et al., 2018; Liaw et al., 1997).

In fungi, PI[4,5]P_2_ can be generated either by a homologue of the mammalian phosphatidylinositol 4-phosphate 5-kinase, Mss-4 (Desrivières et al., 1998; Mähs et al., 2012) or by a homologue of the mammalian PTEN phosphatase (Balla, 2013). Deletion of the PTEN orthologue in *Schizosaccharomyces pombe* resulted in the accumulation of its substrate PI[3,4,5]P_3_, as well as misshapen vacuoles and sensitivity to osmotic stress, while in the budding yeast *Saccharomyces cerevisiae* it caused mislocalisation of dityrosine during sporulation and increased resistance to the phosphoinositide-3 kinase inhibitor, wortmannin (Heymont et al., 2000; Mitra et al., 2004). In *Fusarium graminearum*, mutants lacking *Fgtep1* displayed reduced conidiation, decreased germination in the presence of wortmannin, and reduced virulence on wheat coleoptiles (Zhang et al., 2010). Similarly, in the corn pathogen *Ustilago maydis*, deletion of *Umptn1* caused a decrease in teliospore production, germination and virulence (Vijayakrishnapillai et al., 2018).

Here we investigated the role of PI signalling in two plant-infecting filamentous ascomycetes, *Epichloë festucae* and *Fusarium oxysporum*, that display very different lifestyles. In nature, *E. festucae* grows exclusively within host grasses such as *Lolium perenne*, where it establishes a mutually beneficial symbiotic interaction (Schardl, 2001). This endophyte forms an extensive intercellular hyphal network within the aerial tissues of the plant host as well as epiphyllous hyphae on the plant surface (Becker et al., 2016; Chung and Schardl, 1997; Scott and Schardl, 1993). By contrast, *F. oxysporum* resides mainly in the soil, from where it locates and penetrates the roots of host plants to colonize the xylem vessels and cause devastating vascular wilt disease on more than one hundred crops (Dean et al., 2012; Di Pietro et al., 2003). Intriguingly, both the mutualistic and the pathogenic interaction of these two fungi are exquisitely regulated by conserved signalling pathways, including three mitogen-activated protein kinase (MAPK) pathways, the cell wall integrity (CWI), the invasive growth (IG) and the high-osmolarity glycerol (HOG) pathway, the striatin-interacting phosphatase and kinase (STRIPAK) complex, and reactive oxygen species (ROS) signalling (Becker et al., 2015; Di Pietro et al., 2001; Eaton et al., 2008; Green et al., 2016; Segorbe et al., 2017; Tanaka et al., 2006; Turrà et al., 2015). However, the role of PI signalling during the key steps of symbiosis and root infection is currently unknown. Here we studied the localisation of PIs in *E. festucae* using a purpose-designed suite of PI-binding molecular probes. Furthermore, we analysed the role of two key components in PI signalling, the lipid kinase Mss4 and the lipid phosphatase PTEN, during hyphal growth, development and the *E. festucae*–*L. perenne* interaction, and also tested the role of PTEN in *F. oxysporum* chemotropism.

## Materials and Methods

### Bioinformatic and statistical analyses

A tBLASTn analysis against the *E. festucae* Fl1 and the *F. oxysporum* f. sp. *lycopersici* 4287 genome was performed on the Kentucky Endophyte database (endophyte.uky.edu) (Schardl et al., 2013) and the NCBI BLAST server, respectively, using the Tep1 protein sequence from *S. cerevisiae* (*Sc*Tep1) as a query, to identify EfM3.013870 and FOXG_09154 (hereafter referred to as TepA). Expression analysis of *tepA* in *E. festucae* made use of previous transcriptome datasets (Chujo et al., 2019; Eaton et al., 2015). The protein sequence of TepA was analysed with InterProScan (Jones et al., 2014), and protein alignments were generated using MAFFT (Katoh et al., 2017) and JalView (v1.0). Boxplots were generated online with BoxPlotR (available on http://shiny.chemgrid.org/boxplotr/). Dunn’s test for multiple comparisons, as implemented in the R package dunn.test v1.3.5 (Dinno, 2017) was used to identify significant differences in the number of hyphae per intercellular space between all deletion and over-expression strains. One-way ANOVAs were used to test for differences in plant phenotypes between wild-type (WT) and overexpression strains. In each case, the ANOVA was fitted with R, and a Bonferroni correction was applied to all p-values to account for multiple testing.

### Strains and growth conditions

*Escherichia coli* and *E. festucae* strains were grown at 37°C and 22°C, respectively, as previously described (Green et al., 2017). All strains are listed in Table S1. Conidia were harvested by scrubbing colonies with sterile water followed by filtering through glass wool as previously described (Green et. al., 2017)

The tomato-pathogenic isolate *F. oxysporum* f. sp. *lycopersici* 4287 (FGSC 9935) was used throughout the study. For extraction of DNA and microconidia production, fungal strains were grown at 28°C with shaking at 170 rpm in liquid PDB medium supplemented with the appropriate antibiotics as previously described (Di Pietro and Roncero, 1998).

### Plant growth and inoculation

*E. festucae* strains were inoculated into *L. perenne* seedlings (cv. Samson) and grown as previously described (Becker et al., 2018; Latch and Christensen, 1985).

Tomato root inoculation assays with *F. oxysporum* were performed as previously described (Di Pietro and Roncero, 1998). Roots of 2-week-old *Solanum lycopersicum* seedlings (cv Moneymaker) were immersed for 30 min in a suspension of 5×10^6^ microconidia ml^-1^ and planted in minipots with vermiculite. Plants (ten per treatment) were maintained in a growth chamber (15 h:9 h, light:dark cycle, 28°C). Survival was recorded daily, calculated by the Kaplan-Meier method and compared among groups using the log-rank test.

### DNA analysis, PCR and sequencing

*E. festucae* genomic DNA was isolated from freeze-dried mycelium, as previously described (Byrd et al., 1990). Plasmid DNA was isolated from *E. coli* liquid cultures using the High Pure plasmid isolation kit (Roche, Basel, Switzerland).

PCR amplification of short fragments for screening purposes (<3 kbp) was conducted using OneTaq® (New England Bolabs (NEB), Ipswich, USA) polymerase and of long fragments (>3 kbp) for subsequent cloning procedures using Phusion Polymerase (Thermo Fischer, Waltham, USA). PCR fragments were separated by agarose-gel electrophoresis on a 0.8% gel, and gel extractions were performed with the Wizard SV Gel and PCR clean-up kit (Promega, Madison, USA). Sequencing of plasmids was performed at the Massey Genome Centre. Sequence data was analysed and assembled (ClustalW) using MacVector (v14.5.2).

### Preparation of constructs

All constructs were designed in MacVector (v14.5.2) and assembled via Gibson assembly (Gibson, 2009). A detailed description of all constructs can be found in Methods S1. All primer sequences and generated plasmids can be found in Table S2.

### Transformation of organisms

*E. coli* DH5*α* cells were made chemically competent and transformed as previously described (Hanahan, 1983). *E. festucae* strains were transformed by PEG-mediated protoplast transformation (Young et al., 2005). Protoplasts were generated as previously described (Itoh et al., 1994) and transformed with up to 5 µg of the construct of interest. Following an overnight recovery step on regeneration (RG) medium (Young et al., 2005), protoplasts were overlaid with 0.8% RG agar containing hygromycin B or geneticin to a final concentration of 150 µg/mL or 200 µg/mL, respectively. The gene knockout of *F. oxysporum* protoplasts was generated as previously described (Corral-Ramos et al., 2015).

Western blot *E. festucae* protein extract (30-50 µg), obtained as previously described (Hassing et al., 2020), was resolved on a 7% or 10% SDS acrylamide gel and electrophoretically transferred to nitrocellulose membranes. Membranes were blocked with 2.5% trim milk powder in TBS-T and probed with a primary rabbit anti-GFP antibody (Abcam, Cambridge, UK, ab290, 1:3000 or 1:10000 dilution) or a primary rabbit anti-mCherry antibody (Abcam, ab167453, 1:2000 dilution). A secondary goat anti-rabbit HRP antibody (Abcam, ab6721, 1:10000 dilution) was used for the final antibody incubation, and the blots were developed using Amersham ECL Western blotting solution (GE Healthcare Life Sciences, Malborough, USA).

### qRT-PCR

RNA isolation and qRT-PCR was carried out as previously described using translation elongation factor 2 (*EF-2*) and 40S ribosomal protein S22 (*S22*) as reference genes (Lukito et al., 2015), and primer combinations BH189/BH190 and AC33/AC34 for *tepA* and *mssD*, respectively.

Fungal biomass was quantified using DNA isolated from three biological replicates of pseudostem and blade sections (Liu et al., 2000). Relative biomass was determined by qPCR analysis (SsoFast(tm) EvaGreen®, Bio-Rad) using the ratio of *pacC* (primers YL113F/YL113R) and *hepA* (primers YL120F/YL120R), single copy endophyte genes (Lukito et al., 2015), to *LpCCR1* (primers YL501F/YL501R and YL502F/YL502R), a single copy plant gene (McInnes et al., 2002). Standard curves were generated by the Lightcycler480® (Roche) software (v1.5.0.) for each primer pair using purified PCR products and absolute quantification qPCR performed with two technical replicates per sample. All primer sequences can be found in Table S2.

### Microscopy

Morphology and growth of cultures, grown as previously described (Becker et al., 2015), were analysed using an Olympus IX83 inverted fluorescence microscope. Cell walls and endomembranes were visualised by staining mycelia with Calcofluor white (CFW) (3 µl of a 3 mg/mL solution), and FM4-64 (3 µl of a 16.4 µM solution), respectively.

Pseudostem samples were analysed by confocal laser scanning microscopy after labelling with WGA-AlexaFluor 488 (WGA-AF488, Molecular Probes/Invitrogen) and aniline blue diammonium salt (Sigma, St. Louis, USA) (Becker et al., 2016; Becker et al., 2018) using a Leica SP5DM6000B confocal microscope (488 nm argon and 561 nm DPSS laser, 40*×* oil immersion objective, NA = 1.3) (Leica Microsystems).

### Chemotropic assay

Chemotropic growth of different *F. oxysporum* strains towards tomato root exudate and horseradish peroxidase (HRP; Sigma - Aldrich) was measured using a quantitative plate assay as previously described (Turrà et al., 2015). Microconidia (10^6^) embedded in 0.5% water agar, were incubated for 13 h at 22°C in the presence of a chemoattractant gradient, and the direction of germ tubes relative to a central scoring line was determined using an Olympus binocular microscope at 9200*×* magnification. For each sample, five independent batches of cells (n = 100 cells per batch) were scored. The chemotropic index was measured as previously described (Turrà et al., 2015). All experiments were performed at least three times with similar results. Statistical analysis was conducted using the *t*-test for unequal variances, also referred to as Welch’s test.

## Results

### Distribution of phosphoinositides in *Epichloë festucae* membranes

To carry out a comprehensive microscopic analysis of PIs in *E. festucae*, fluorescent molecular probes for the PI species PI[3]P, PI[4]P, PI[4,5]P_2_, PI[3,4]P_2_, PI[3,5]P_2_ and PI[3,4,5]P_3_ were generated. Each probe consists of a characterised mammalian lipid-binding domain specific to one of the respective PIs translationally fused to a C-terminal eGFP or an N-terminal mCherry (Table 1). Codon-optimised constructs were expressed under the control of the *Aspergillus nidulans gpdA* promoter, which drives high level protein expression in *E. festucae* (Hassing et al., 2020), and transformed into *E. festucae* Fl1 protoplasts. Construct integration was verified by PCR and correct expression of the molecular probes was subsequently confirmed by western blot analysis (Fig. S1). Because strain Fl1 produces very few asexual spores, the constructs were also transformed into *E. festucae* strain E2368, a prolific producer of conidiospores (Wilkinson et al., 2000).

**Table 1:** Molecular probes generated for this study.

| Mammalian protein | Accession | Lipid-binding domain | Lipid target | Reference |
| --- | --- | --- | --- | --- |
| Hepatocyte growth factor-regulated tyrosine kinase substrate (HGS) | NP_001152800 | FYVE (aa 15-222) | PI[3]P | (Vermeer Joop et al., 2006) |
| Tandem PH-domain-containing protein-2 (Plekha2) | NP_112547 | PH (aa 195-304) | PI[3,4]P<sub>2</sub> | (Dowler et al., 2000) |
| Mucolipin 1 (TRPML1) | NP_444407.1 | tandem repeat of lipid binding domain (aa 1-68) | PI[3,5]P<sub>2</sub> | (Li et al., 2013) |
| Bruton's tyrosine kinase (BTK) | NP_038510 | PH (aa 3-169) | PI[3,4,5]P<sub>3</sub> | (Rameh et al., 1997) |
| Four-phosphate-adaptor protein 1 (Plekha3) | NP_112546 | PH (aa 1-96) | PI[4]P | (Vermeer Joop et al., 2006) |
| Phospholipase C-δ1 (PLC) | NP_062650 | PH (aa 13-139) | PI[4,5]P<sub>2</sub> | (Van Leeuwen et al., 2007b) |

The localisation of PI[3,4]P_2_ and PI[3,4,5]P_3_ probes in *E. festucae* hyphae grown in axenic culture or *in planta* was comparable to that of free eGFP and mCherry proteins (Fig. S2). This suggests that *E. festucae* lacks PI[3,4]P_2_ and PI[3,4,5]P_3_, or that these lipid species are present at concentrations below the detection limit. By contrast, we found that the PI[3]P probe localised to mobile vesicles in conidia and hyphal tips, as well as to the periphery of large oval organelles, presumably vacuoles, present only in mature hyphae (Fig. 1). Localisation of the PI[3]P probe to endocytic vesicles and the vacuolar membrane was confirmed by co-localisation with the membrane marker FM4-64 (Fig. 2). During symbiosis with the plant host, the PI[3]P probes showed a similar distribution to free eGFP or mCherry, localising uniformly throughout the fungal hyphae or in a punctate pattern (Fig. 3). Probes designed for the localisation of PI[3,5]P_2_ showed a divergent localisation pattern in hyphal tips and mature hyphae. While the probe translationally fused to eGFP at the C-terminus localised to mobile, vesicle-like structures in hyphal tips and to the periphery of large, oval organelles, presumably vacuoles, in mature hyphae, the probe with the N-terminal mCherry fusion localised to the cytoplasm and within these round organelles. Analysis of the fusion proteins by western blot suggests the mCherry fusion proteins are more susceptible to degradation (Figs 1, S1).

**Figure 1.**
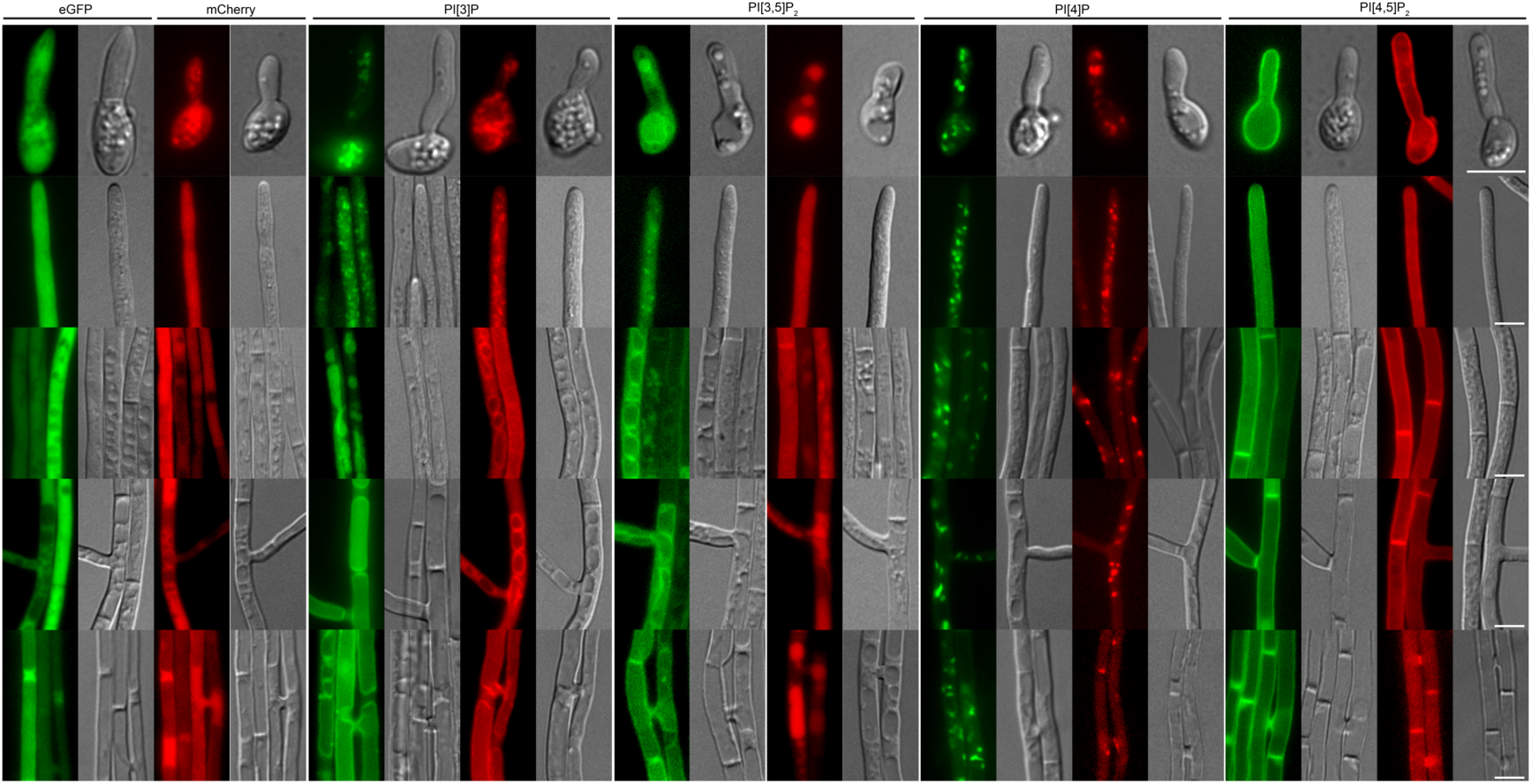
Localisation of the PI[3]P-, PI[3,5]P_2_-, PI[4,]P-, and PI[4,5]P_2_-molecular probes in *Epichloë festucae* hyphae grown in axenic culture. Strains were grown on 1.5% H_2_O agar for 5 d before examination using a fluorescence microscope. Images shown are representative of all strains analysed and show localisation of the molecular probes in hyphae of different ages. Images of spores were acquired from transformed *E. festucae* E2368 strains (with the exception of ML1 molecular probes), whereas remaining ones are from transformed Fl1 (with the exception of PI[3,5]P_2_ molecular probes). Cytosolic eGFP (pCE25): #T5 (E2368), #T8 (Fl1); cytosolic mCherry (pCE126): #T7 (E2368), #T2 (Fl1), HGS-eGFP (PI[3]P, pCE106): #T7 (E2368), #T9 (Fl1); mCherry-HGS (PI[3]P, pCE111): #T29 (E2368), #T37 (Fl1); ML1-eGFP (PI[3,5]P_2_, pBH87): #T5 (spores and axenic culture, Fl1); mCherry-ML1 (PI[3,5]P_2_, pBH88): #T12 (spores), #T6 (axenic culture, Fl1); Plekha3-eGFP (PI[4]P, pCE109): #T2 (E2368), #T3 (Fl1); mCherry-Plekha3 (PI[4]P, pCE114): #T16 (E2368), #T25 (Fl1); PLC-eGFP (PI[4,5]P_2_, pCE105): #T10 (E2368), #T3 (Fl1); mCherry-PLC (PI[4,5]P_2_, pCE110): #T5 (E2368), #T16 (Fl1); Bar = 5 µm.

**Figure 2.**
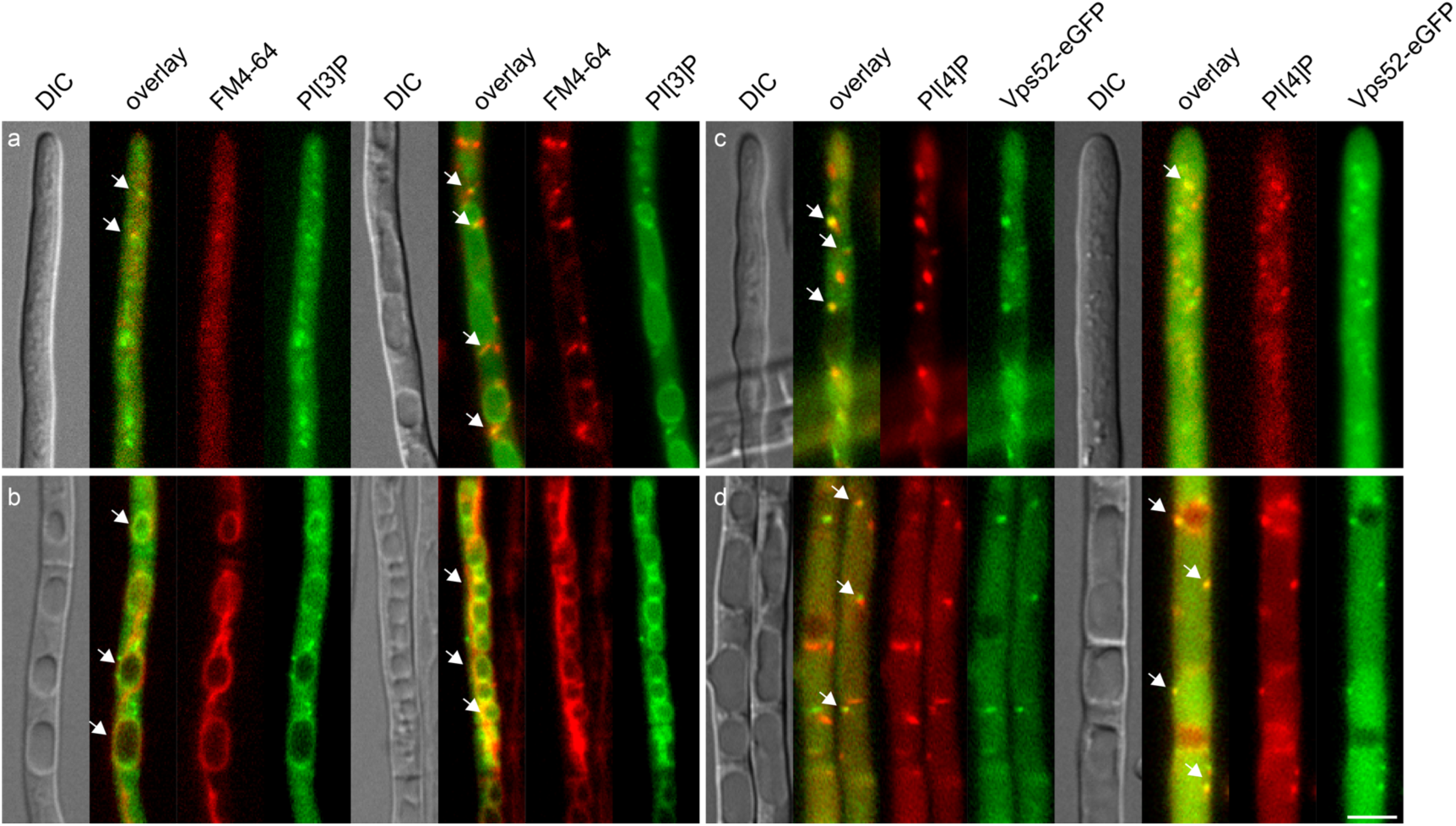
Co-Localisation with compartment-specific fluorescent markers of the PI[3]P- and PI[4]P-molecular probes in *Epichloë festucae* hyphae grown in axenic culture. Strains were grown on 1.5% H_2_O agar for 5 d before examination using a fluorescence microscope. Images shown are representative of all strains analysed and show localisation of the molecular probes in hyphae of different ages. (a, b) Co-localisation of the PI[3]P molecular probe with the membrane marker FM4-64. In (a) FM4-64 was added and cultures were analysed immediately to observe endocytic vesicles, while in (b) the stain was added and the cultures were incubated 16 h before analysis, resulting in the staining of vacuolar membranes. Images shown are from the strains HGS-eGFP (PI[3]P, pCE106) #T9 and #T24. (c and d) Co-localisation of the PI[4]P-molecular probe with Vps52-eGFP (pKG55). In (c) hyphal tips are shown, where co-localisation was frequently difficult to observe due to cytoplasmic movement, while (d) shows more mature hyphae. Images are from the strain mCherry-Plekha3 (PI[4]P, pCE114) #T25 transformed with the plasmid pKG55 (Vps52-eGFP) resulting in the strains #T35 and #T43. White arrows show areas of co-localisation. Bar = 5 µm.

**Figure 3.**
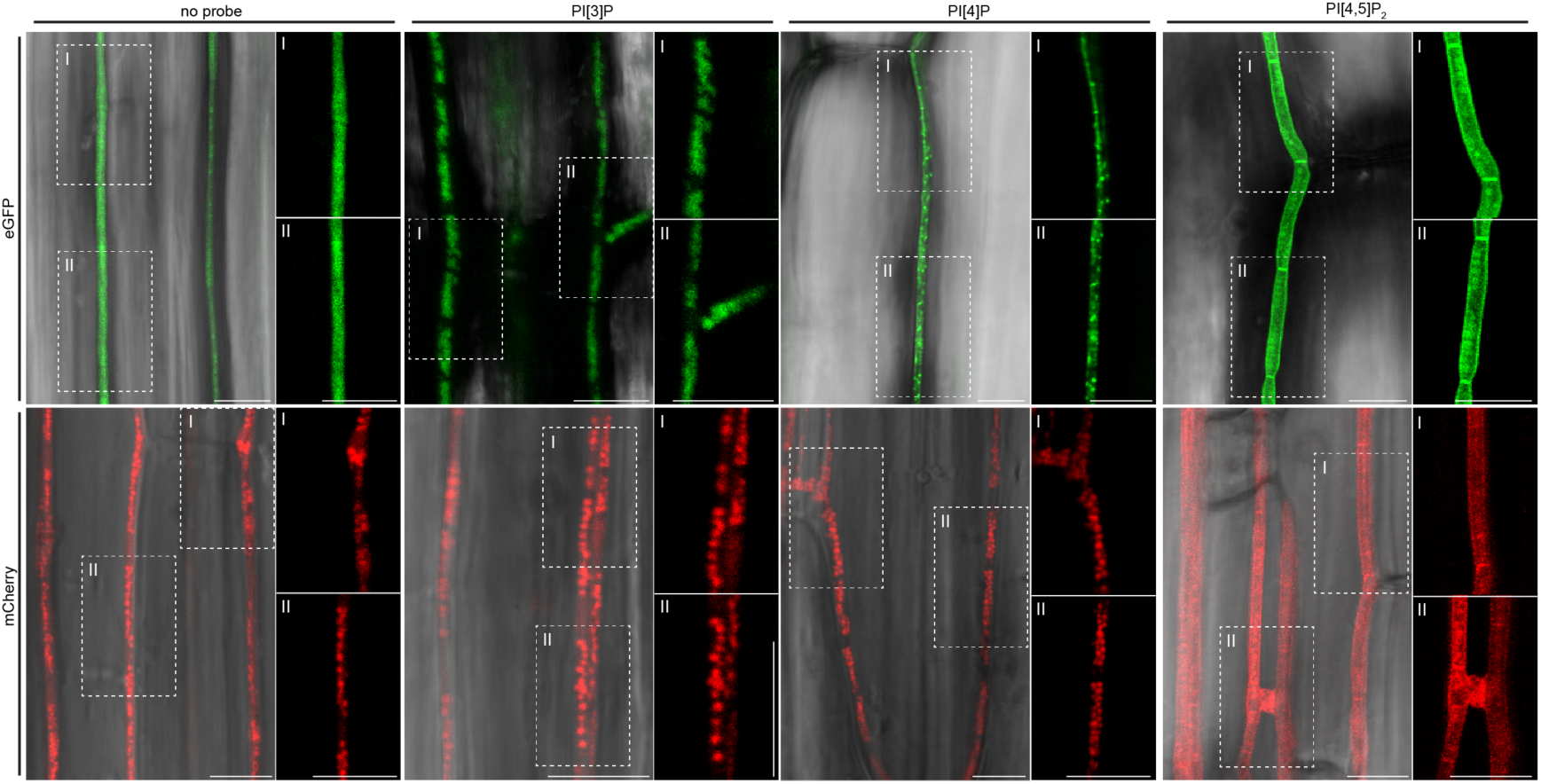
Localisation of the PI[3]P-, PI[4]P- and PI[4,5]P_2_- molecular probes in *Epichloë festucae* during interaction with *Lolium perenne*. Confocal scanning laser microscopy images of *E. festucae* strains constitutively expressing eGFP, mCherry or the PI molecular probes *in symbio* with *L. perenne*. Samples were freshly harvested from the innermost layer of the pseudostem at 9-14 weeks post-inoculation and mounted in deionised water for analysis. Images labeled with (I) or (II) show enlarged images of the region indicated in the neighbouring image. Cytosolic eGFP (pCE25): #T8; cytosolic mCherry (pCE126): #T2; HGS-eGFP (PI[3]P, pCE106): #Tx; mCherry-HGS (PI[3]P, pCE111): #Tx; Plekha3-eGFP (PI[4]P, pCE109): #T3; mCherry-Plekha3 (PI[4]P, pCE114): #T25; PLC-eGFP (PI[4,5]P_2_, pCE105): #T6; mCherry-PLC (PI[4,5]P_2_, pCE110): #T10; Bar =10 µm.

Both probes occasionally localised to septa (Fig. 1). The PI[4]P probe localised to small mobile vesicles in conidia, hyphal tips and mature hyphae, during growth in axenic culture and *in planta* (Figs 1, 3). In older hyphae, the cytoplasmic background signal was frequently increased, suggesting a lack of PI[4]P or a higher rate of probe degradation in these hyphae (Fig. 1). Co-localisation with Vps52-eGFP, a marker protein for late Golgi vesicles, confirmed that some of the PI[4]P-containing vesicles were late Golgi vesicles (Fig. 2). Interestingly, the PI[4,5]P_2_ probe localised to the periphery and septa of hyphae of all developmental stages, both in culture and *in planta* (Figs 1, 3). Strikingly, the distribution at the hyphal apex was highly asymmetric, with a reduced signal at the tip and higher fluorescence in the subapical region (Figs 1, S3).

### *Epichloë festucae* encodes a phosphatidylinositol 4-phosphate 5-kinase and a homologue of the mammalian lipid phosphatase and tumour suppressor protein PTEN

We next analysed the role of lipid phosphatases and kinases on hyphal growth and localisation of PI molecular probes in *E. festucae*. The homologue of the *S. cerevisiae* Mss4 phosphatidylinositol 4-phosphate 5-kinase was identified from a tBLASTn analysis of the *E. festucae* Fl1 genome, using the protein sequence of Mss4 (YDR208W) as the query. This search identified a gene (EfM3.031950) encoding a protein of 963 aa (*E*-value 0), hereafter designated as MssD. MssD is the *E. festucae* homologue of the mammalian phosphatidylinositol 4-phosphate 5-kinase type-1 alpha/beta, and contains a highly conserved phosphatidylinositol-4-phosphate 5-kinase domain (IPR023610) (Jones et al., 2014) (Fig. 4). Embedded in this domain is a monopartite nuclear localisation signal, as described previously for *S. cerevisiae* Mss4 (Audhya and Emr, 2003).

**Figure 4.**
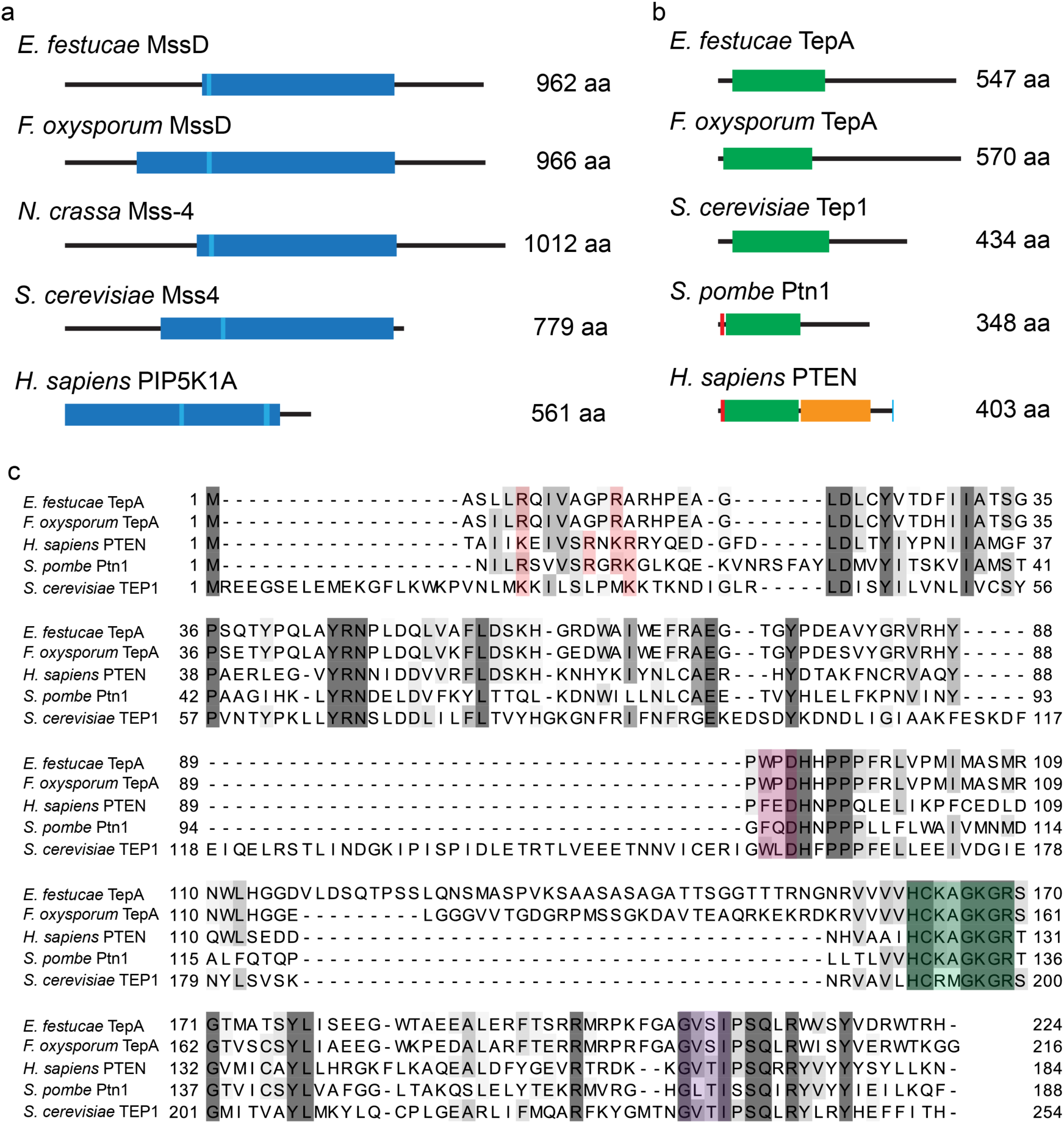
Domain structure of *Epichloë festucae* MssD and TepA, and comparison to homologues from other fungi and humans. (a) Schematic representation of *E. festucae* MssD, *F. oxysporum* MssD (XP_018251113.1), *Neurospora crassa* Mss-4 (NCU02295), *S. cerevisiae* Mss4 (YDR208W), and human PIK5P1A (Q99755). Dark blue box: phosphatidylinositol-4-phosphate 5-kinase domain (IPR023610), light blue box: Nuclear localisation signal (cNLS Mapper, (Kosugi et al., 2008; Kosugi et al., 2009a; Kosugi et al., 2009b)), human PIK5P1A contains a bipartite NLS. Size of the protein indicated in number of aa. (b) Schematic representation of *E. festucae* TepA, *Schizosaccharomyces pombe* Ptn1 (CAA22831.1), *Saccharomyces cerevisiae* Tep1 (P53916.1), *Fusarium oxysporum* TepA (XP_018246213.1) and *Homo sapiens* PTEN (AAD13528.1). Green box: tensin-type phosphatase domain, red box: PI[4,5]P_2_-binding domain, orange box: C2 domain, blue box: PDZ binding motif. Size of the protein indicated in number of aa. (c) Amino acid sequence alignment of the tensin-type phosphatase domain of TepA, *Sp*Ptn1, *Sc*TEP1, *Fo*TepA and *Hs*PTEN; red highlights: conserved aa of the PI[4,5]P_2_-binding domain; light purple box: WPD-loop; green box: P-loop (active site); dark purple box: Ti-loop. Sequences were annotated according to InterProScan (Jones et al., 2014) and by comparison of the domains identified by (Lee et al., 1999). Sequences are shaded in a grey scale based on their percentage identity.

We next identified the *E. festucae* homologue of the mammalian tumour suppressor protein PTEN by performing a tBLASTn analysis against the Fl1 genome using *S. cerevisiae* Tep1 (P53916.1) as a query. This search returned one putative homologue, EfM3.013870, (*E*-value 5e-13), designated as *tepA*, which encodes a predicted protein of 547 aa. Analysis of TepA using InterProScan (Jones et al., 2014) revealed the presence of a dual-specificity phosphatase activity domain (153–202 aa) embedded in the tensin-type phosphatase domain, characteristic of PTEN-type proteins from fungi to humans. All conserved aa residues of the catalytic P-, WPD- and TI-loops were present in TepA (Fig. 4) (Lee et al., 1999), while the anionic lipid-binding C2 domain and the PDZ (PSD-95, Discs-large, ZO-1)-binding motif were lacking in TepA, as described for all fungal homologues analysed to date (Cid et al., 2008). Two of the four conserved aa of the N-terminal PI[4,5]P_2_-binding domain (PBD; K/R-X_4_-K/R-X-K/R-K/R), required for activity and localisation of PTEN, were absent in TepA as previously reported for *Fg*Tep1 (Campbell et al., 2003; Walker et al., 2004) (Fig. 4). Protein sequence alignment of the tensin-type phosphatase domain of PTEN, *Fg*Tep1, TepA, *Sp*Ptn1 and *Sc*Tep1 demonstrated an *S. cerevisiae*-specific insertion in *Sc*Tep1, as well as an apparent filamentous fungus-specific insertion in TepA and *Fg*Tep1 (Cid et al., 2008).

Analysis of *tepA* and *mssD* gene expression, making use of available transcriptome datasets (Chujo et al., 2019; Eaton et al., 2015; Hassing et al., 2019), revealed an intermediate level of expression in culture (20.76 RPKM and 34.57 RPKM, respectively) but no differential expression *in planta* or in any of the symbiotic mutants analysed to date (*ΔsakA, ΔnoxA, ΔproA*, or *ΔhepA*).

### *Epichloë festucae* MssD and TepA exhibit distinct localisation patterns

Mammalian PTEN has been described to bind to the PM over a very short period (hundredths of a millisecond) to dephosphorylate PI[3,4,5]P_3_ before dissociating again to the cytoplasm (Vazquez et al., 2006). By contrast, *Neurospora crassa* Mss-4 associates stably with the PM in the subapical region and in filamentous structures (Mähs et al., 2012). To analyse the localisation of TepA in *E. festucae*, we designed N-terminal mCherry (pBH41) and C-terminal eGFP (pBH42) fusion constructs based on the genomic DNA sequence of *tepA*. For MssD, we designed an N-terminal eGFP fusion construct (pBH70), analogous to that used for successful localisation of *Nc*Mss-4 (Mähs et al., 2012). These constructs were transformed into wild-type (WT) protoplasts and expression of the complete fusion proteins was verified via western blot (Fig. S4). The eGFP-MssD fusion protein localised to the PM and septa of hyphae of all ages (Fig. 5). Localisation to the PM was irregular and punctate and the fluorescence signal at the hyphal tips was very weak, in contrast to the extended PM localisation of the PI[4,5]P_2_ molecular probe (Fig. 1). In mature hyphae, the signal was frequently observed in the membrane of small round organelles. For the TepA-mCherry and the eGFP-TepA fusion proteins, slight differences in the localisation were observed. The eGFP-TepA fusion protein localised in the cytoplasm of hyphal tips and young hyphae, similar to constitutively expressed eGFP and mCherry. By contrast, the TepA-mCherry localised to the cytoplasm as well as to septa in hyphae of all ages (Fig. 5).

**Figure 5.**
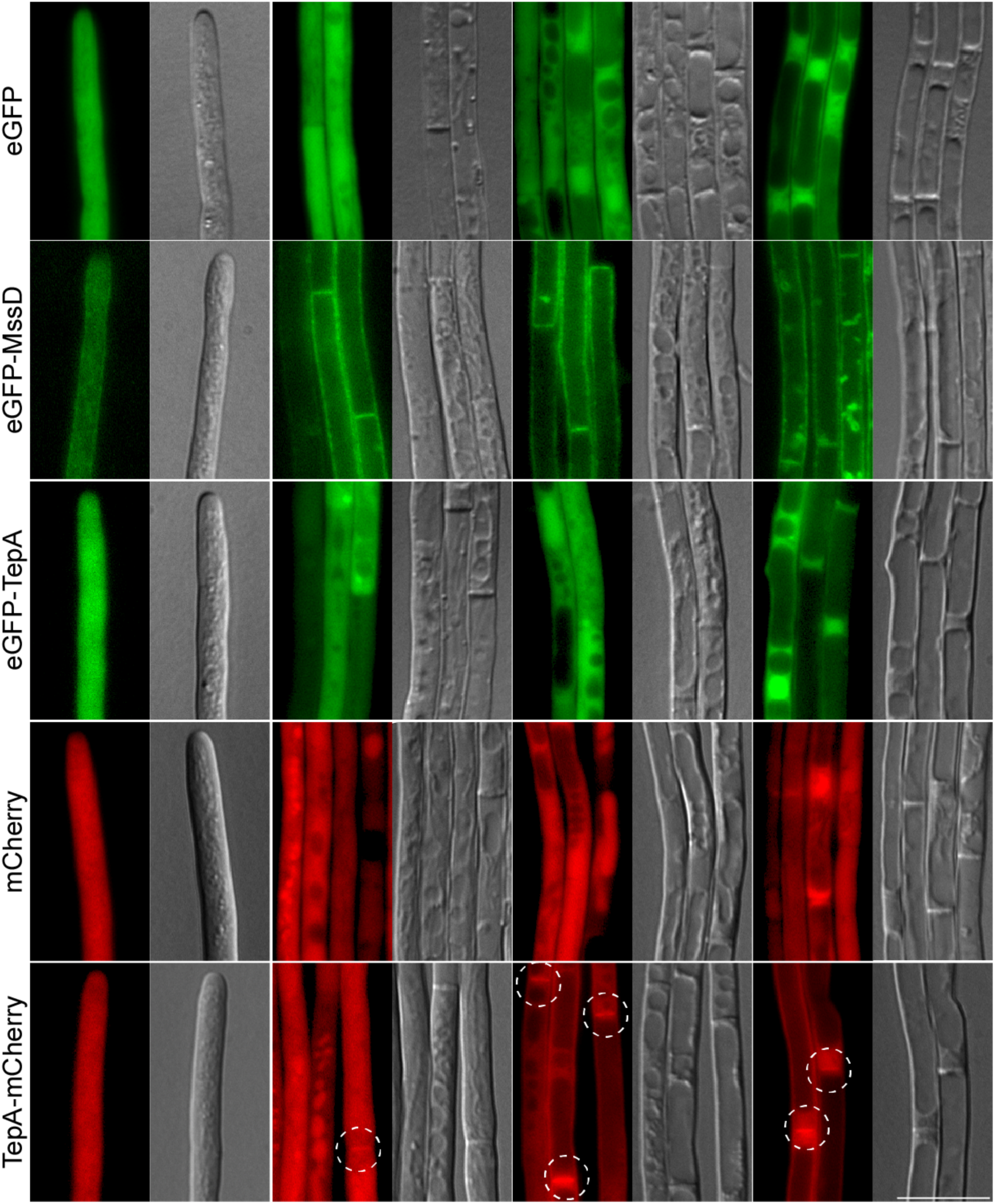
Localisation of TepA and MssD during axenic growth of *Epichloë festucae*. Strains were grown on 1.5% H_2_O agar for 5 d before examination using a fluorescence microscope. The images shown of wild-type (WT) expressing eGFP (pBH28, #T6), eGFP-MssD (#T8), eGFP-TepA (pBH41, #T4), mCherry (pCE126, #T2) and TepA-mCherry (pBH42, #T2) are representative of all strains analysed. As indicated, hyphae of different ages were analysed. White circles: localisation of the fusion protein at septa. Bar = 10 µm.

### Loss of *tepA* leads to accumulation of PI[3,4,5]P_3_ in hyphal septa

To test if TepA and MssD affect the concentration and localisation of different PIs, we generated *mssD* and *tepA* deletion and overexpression strains. Gene replacement by homologous recombination yielded three *E. festucae tepA* deletion strains named #T80, #T87 and #T102 (Fig. S5). By contrast, no *mssD* deletion strains were identified after screening ∼200 transformants, suggesting that *mssD* is essential in *E. festucae*, as previously described in *S. cerevisiae* and *N. crassa* (Mähs et al., 2012; Yoshida et al., 1994).

MssD and TepA overexpression strains were generated by transforming WT protoplasts with plasmids containing the *mssD* (pCE101) or *tepA* (pCE122) gene under the control of the *A. nidulans gpdA* promoter. The transcript levels of *mssD* and *tepA* in a number of transformants were determined by RT-qPCR, and strains that showed increased transcript levels compared to the WT were used for subsequent studies: *mssD* OE: #T17, #T20, #T53 and *tepA* OE: #T4, #T7, #T20 (Fig. S6). Interestingly, all of the *mssD* overexpression (OE) transformants analysed showed relatively low levels of *mssD* overexpression (Fig. S6), suggesting that a high expression level of *mssD* could be detrimental to *E. festucae*.

We then tested whether the overexpression of *mssD* or *tepA*, or the deletion of *tepA* affected the concentration and localisation of different PIs by transforming these constructs into protoplasts of WT, the *tepA* deletion strains #T87 and #T102, the *tepA* overexpression strain #T7 and the *mssD* overexpression strain #T17, which showed the highest levels of expression in culture. Using the PI[4,5]P_2_ or the PI[3,4,5]P_3_ molecular probes, a signal pattern similar to that of WT hyphae was observed in the different mutant strains (Figs 1, 6). However, the *ΔtepA* mutants showed an accumulation of PI[3,4,5]P_3_ at septa in hyphae of all ages (Fig. 6), supporting the hypothesis that *E. festucae* TepA is a functional phosphatidylinositol 3,4,5-trisphosphate 3-phosphatase.

**Figure 6.**
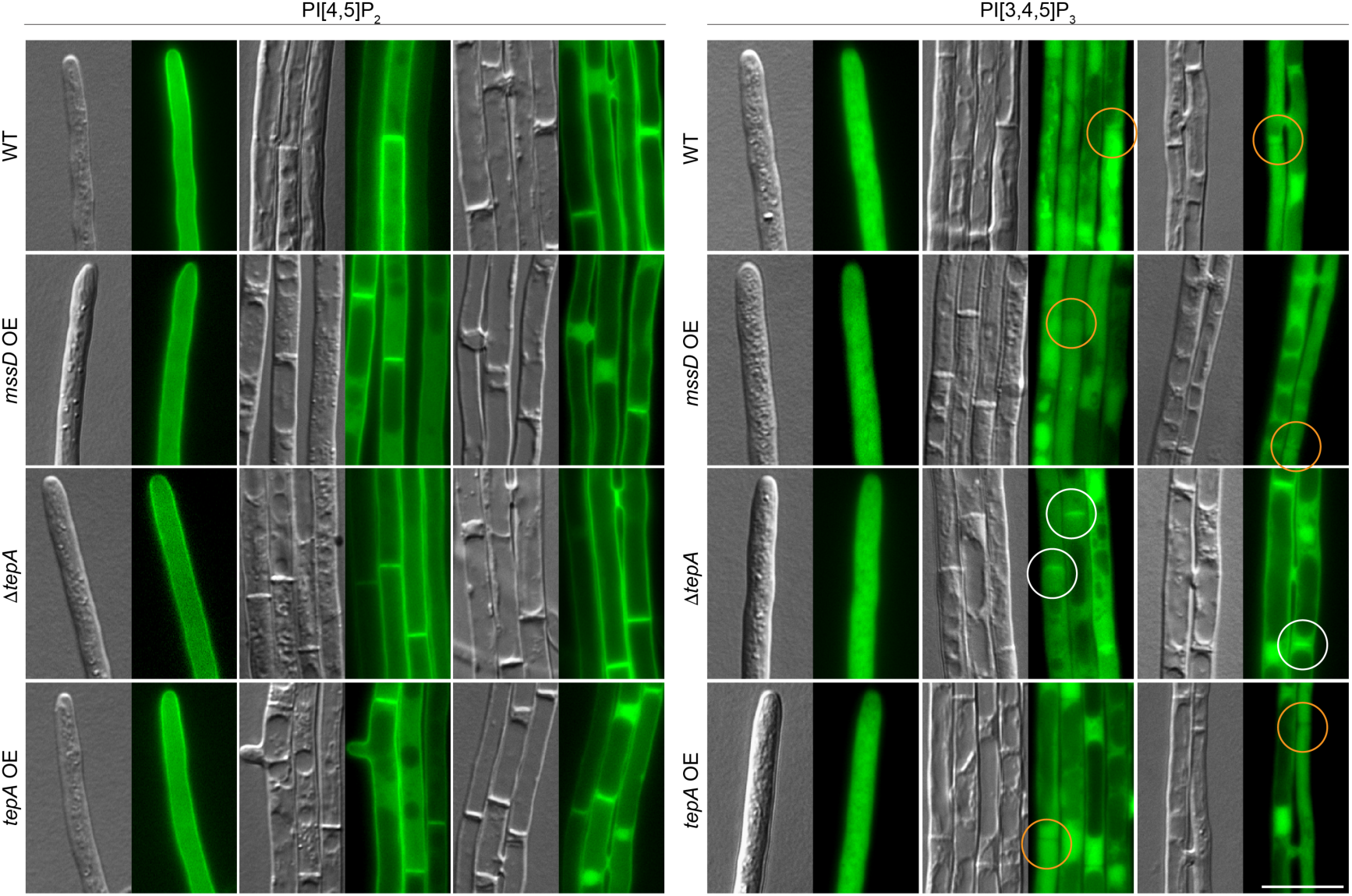
Localisation of PI[4,5]P_2_ and PI[3,4,5]P_3_ in *Epichloë festucae* wild-type, *tepA* and *mssD* mutant strains. The indicated strains were incubated on 1.5% H_2_O agar for approx. 5 d before analysis by fluorescence microscopy. eGFP-based Molecular probe constructs were transformed into WT, the *tepA* deletion strain #T87, the *tepA* overexpression strain #T7 and the *mssD* overexpression strain #T17. Multiple transformants were analysed and images shown are representative of all transformants analysed. (a) Localisation of the PI[4,5]P_2_ molecular probe (pCE107) in WT (#T3), *mssD* overexpression (#T8), *tepA* overexpression (#T6) and *tepA* deletion (#TN2) strains. (b) Localisation of the PI[3,4,5]P_3_ molecular probe (pCE105) in WT (#T9), *mssD* overexpression (#T8), *tepA* overexpression (#T1) and *tepA* deletion (#T15) strains. White circles: localisation of the molecular probe at septa. Orange circles: No localisation of the molecular probe at septa. Bar = 10 µm.

### Overexpression of *Epichloë festucae mssD* or *tepA*, or deletion of *tepA* does not affect hyphal growth in culture

To analyse the role of MssD and TepA in *E. festucae*, growth of WT and mutant strains were compared in axenic culture. The growth rate and morphology of the *mssD* overexpression strains and the *tepA* deletion and overexpression strains were indistinguishable from the WT (Fig. S7). Similar to the WT, these strains showed smooth hyphae of a uniform diameter, formation of hyphal bundles, hyphal coils and hyphal cell-to-cell fusions (Fig. S7). Cell wall staining of WT and mutant hyphae with the chitin-binding dye Calcofluor white (CFW) resulted in a similar even pattern of fluorescence (Fig. S7). We conclude that overexpression of *mssD* or *tepA*, or deletion of *tepA* have no detectable effect on the growth of *E. festucae* in axenic culture.

### Overexpression of *Epichloë festucae mssD* or *tepA* does not affect fungal growth *in planta*

In previous studies, deletion of the PTEN homologues in *F. graminearum* and *U. maydis* resulted in a decrease in virulence on wheat and maize, respectively (Vijayakrishnapillai et al., 2018; Zhang et al., 2010). Here we found that *L. perenne* plants infected with the *tepA* deletion mutant of *E. festucae* and scored 10 weeks post-inoculation, were not consistently significantly different from WT-infected plants in leaf length and tiller number (Figs 7, S8). Similar results were observed for plants infected with the *tepA* overexpression strains. Interestingly, plants infected with *mssD* overexpression strains #T20 and #T53 had significantly longer tillers than those infected with the WT (Figs 7, S8). This phenotype was previously correlated with a decreased fungal biomass *in planta* (Kayano et al., 2018). We therefore measured the fungal biomass in plants using qPCR and found that the fungal biomass was significantly reduced in plants infected with the *mssD* overexpression strain #T20 and appeared to be reduced (albeit not always significantly) in plants infected with the overexpression strain #T53 (Fig. S9).

**Figure 7.**
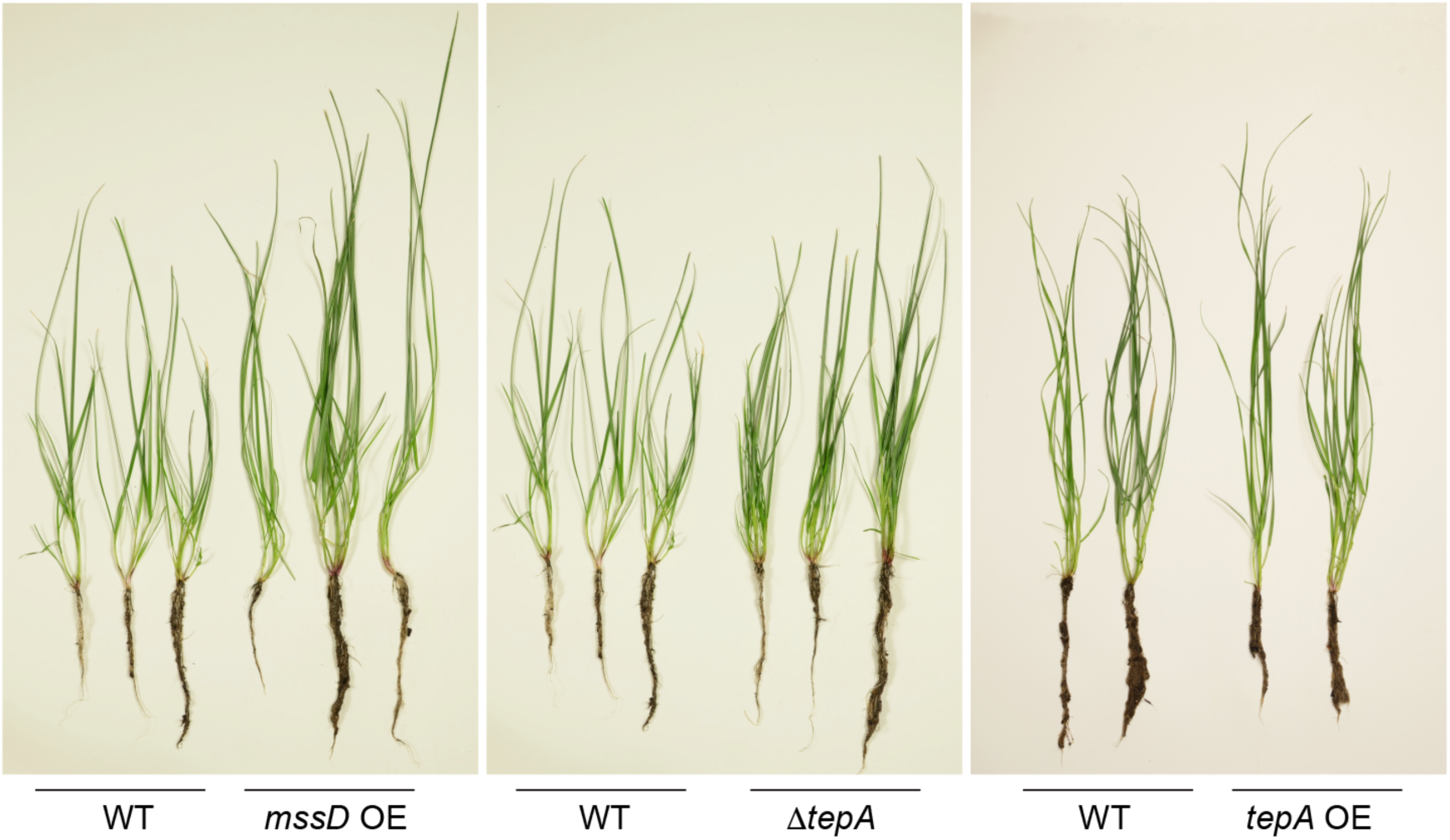
Phenotype of *Lolium perenne* plants infected with *Epichloë festucae* wild-type or different deletion or overexpression strains. Plants infected with the wild-type (WT), *mssD* overexpression, and *tepA* deletion or overexpression strains were scored 10 weeks post-inoculation. Number of infected plants: WT (23), *mssD* overexpression strains <u>#T17</u>, #T20, #T53 (18/15/6), *tepA* deletion strains #T80, <u>#T87</u>, #T102 (31/32/18), WT (5), *tepA* overexpression strains #T4, <u>#T7</u>, #T20: (5/3/2). Plants infected with <u>underlined strains</u> are shown in the figure.

To visualize fungal hyphae in the plant tissue, longitudinal sections of the pseudostem of infected plants were infiltrated with the chitin-specific probe WGA-AF488 and the *β*-glucan-binding dye aniline blue. The growth pattern of the different mutant strains was similar to that of the WT, showing one or two hyphae between neighbouring plant cells, which branch and fuse with neighbouring hyphae to give rise to a hyphal network throughout the plant (Fig. S10).

### *Fusarium oxysporum* TepA is required for chemotropic hyphal growth towards tomato roots

Next we tested the role of TepA in the fungal pathogen *F. oxysporum* by generating targeted deletion mutants in the *tepA* ortholog *FOXG_09154*. Southern blot analysis identified a number of transformants in which the wild-type gene had been replaced with the null allele (Fig. S11). Colony growth and conidiation of these mutants was indistinguishable from the WT. Moreover, the mutants did not significantly differ from the WT in causing mortality in tomato plants in a root inoculation assay (Fig. S12). These results suggest that TepA is not required for hyphal growth, development and virulence of *F. oxysporum*.

PI[3,4,5]P_3_ signalling was shown to be required for cell polarity and chemotaxis in mammalian cells and in the slime mould, *Dictyostelium* (Balla, 2013; Heit et al., 2008; Iijima and Devreotes, 2002; Iijima et al., 2004; Weiger and Parent, 2012). Previously work showed that *F. oxysporum* hyphae are chemotropically attracted to tomato roots by secreted plant peroxidases (Turrà *et al*., 2015). Here we found that the *tepA* deletion mutants were impaired in directional growth towards tomato root exudate and commercial horseradish peroxidase (Fig. 8). This suggests that PI[3,4,5]P_3_ signalling is required for chemotropism of *F. oxysporum* towards plant roots.

**Figure 8.**
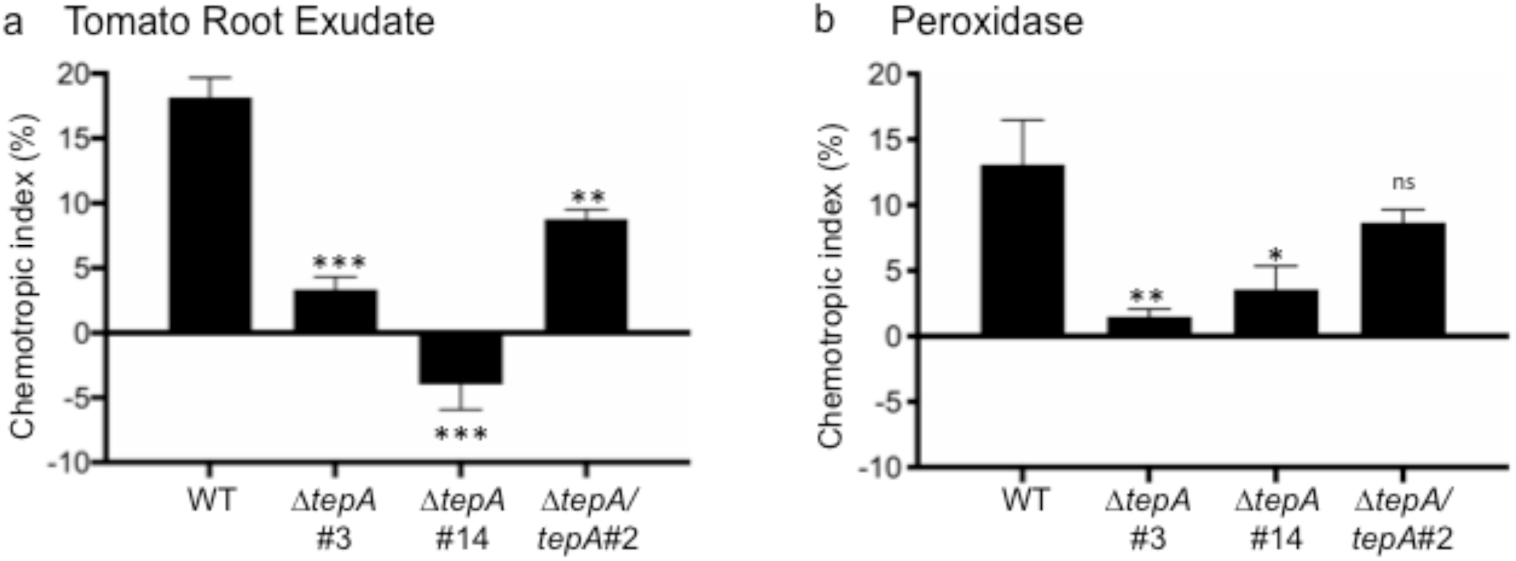
*Fusarium oxysporum* TepA is required for hyphal chemotropism towards tomato roots. Microconidia of the *F. oxysporum* wild type strain, two knockout mutants and a complemented strain were embedded in water agar and incubated 13 h at room temperature in the presence of a gradient of tomato root exudate (a) or 4 µM horseradish peroxidase (HRP) (b). The direction of germ tubes relative to a central scoring line was determined in an Olympus binocular microscope. Calculation of chemotropic index was done as described (Turrà et al., 2015). \**p*<0.05; **, *p*<0.01; ***, *p*<0.001, *versus* wild-type (WT) strain. Data presented are the mean from at least three independent experiments. n = 500 germ per experiment. Error bars, SD.

## Discussion

Phosphoinositides (PI) have been implicated in many key cellular processes including cytoskeleton rearrangement, establishment of cell polarity, regulation of ion channels and the control of cellular proliferation and cancer (De Craene et al., 2017). In this study, we reveal for the first time the localisation of the PI species PI[3]P, PI[3,5]P_2_, PI[4]P and PI[4,5]P_2_ in a filamentous ascomycete, and provide evidence that this species is able to produce PI[3,4,5]P_3_. While we found that the homologues of the mammalian PTEN protein are not required in the mutualistic interaction between *E. festucae* and *L. perenne* and the pathogenic interaction between *F. oxysporum* and tomato, we identified PTEN to be essential for the ability of fungal hyphae to undergo chemotrophic growth towards plant roots.

The molecular probe for PI[3]P (Vermeer et al., 2006) displayed two distinct localisation patterns in cells of axenic cultures: vesicle-like structures, some of which are endocytic vesicles, and the periphery of vacuoles. Similar observations were made in mammalian and yeast cells, where PI[3]P was identified in early endosomes and in internal vesicles of both multivesicular endosomes and vacuoles (Gillooly et al., 2000; Marat and Haucke, 2016), fulfilling key roles in membrane trafficking and vesicle fusion throughout the endosomal system (Balla, 2013; Marat and Haucke, 2016).

The PI[3,5]P_2_ probe (Li et al., 2013) fused to eGFP at the C-terminus localised to vacuolar membranes in mature hyphae and septa, and to mobile vesicles in hyphal tips. PI[3,5]P_2_ is generated and regulated by a vacuolar membrane protein complex in yeast and mammals, and localises to the vacuolar/lysosomal membranes and endosomes, where it is involved in late endosomal protein sorting and autophagy (De Craene et al., 2017; McCartney et al., 2014). Whether the mobile vesicles correspond to endosomes remains to be seen.

The molecular probe for PI[4]P (Vermeer et al., 2009) localised to highly mobile vesicle-like structures throughout hyphae in culture and *in planta*, some of which were Golgi-derived vesicles. In yeast and mammals, PI[4]P localizes in two distinct pools, at the Golgi and the PM, which are maintained through the presence of different PI4Ks (De Craene et al., 2017; De Matteis et al., 2013). In yeast, Golgi localised PI[4]P is crucial for vesicle formation in anterograde and retrograde transport, while the PM pool is required for targeting specific proteins to the PM, resulting in distinct functions in actin cytoskeleton organization, cell wall integrity, and receptor-mediated endocytosis (Audhya and Emr, 2003; Audhya et al., 2000; Baird et al., 2008; Ghugtyal et al., 2015; Hammond et al., 2012; Yamamoto et al., 2018).

The PI[4,5]P_2_ molecular probe (Van Leeuwen et al., 2007a) localised to the PM and septa of hyphae in axenic culture and *in planta*. Indeed, PI[4,5]P_2_ is the major phosphoinositide at the inner leaflet of the PM across kingdoms, where it serves as a precursor for other signalling molecules and fulfils roles in cell polarity, endo- and exocytosis, actin cytoskeleton rearrangement and activation of transporters (De Craene et al., 2017; Di Paolo and De Camilli, 2006). Strikingly, the PI[4,5]P_2_ molecular probe had an asymmetric localisation in vegetatively growing hyphal tips, where there was little signal at the hyphal apex, but a strong signal in the sub-apical region. An asymmetric distribution of PI[4,5]P_2_ at sites of polar growth has been observed in many organisms including tobacco pollen tubes and in *N. crassa*, and is required for the yeast to hyphal transition of *Candida albicans* (Ischebeck et al., 2008; Mähs et al., 2012; Vernay et al., 2012). In contrast to this study, the fluorescent signal for PI[4,5]P_2_ in *C. albicans, S. cerevisiae and N. crassa* was strongest at the hyphal apex (Guillas et al., 2013; Mähs et al., 2012; Vernay et al., 2012). Interestingly, the subapical concentration of this molecular probe coincided with the so-called endocytic collar, a region with increased endocytic activity (Echauri-Espinosa et al., 2012; Taheri-Talesh et al., 2008). While the cause of this asymmetry is unknown, there are several possibilities such as localised activity of phospholipase C (Dowd et al., 2006; Helling et al., 2006) or TepA or an asymmetric localisation of MssD.

While MssD and its homologues are relatively conserved, the *E. festucae* TepA shows differences to human PTEN regarding different lipid binding motifs. Studies suggest that the PI[4,5]P_2_-binding domain, the catalytic site and the C2 domain are required for PTEN lipid binding, while the C-terminal tail is involved in regulating protein activity (Lee et al., 1999; Nguyen et al., 2014; Vazquez et al., 2006; Walker et al., 2004). Therefore differences in the ability of the *E. festucae* protein to bind, and to be recruited, to membranes are likely.

TepA localised to the cytosol as well as to septa of hyphae grown in axenic culture, a result similar to what has been observed for the PTEN homologue in *S. pombe* (Mitra et al., 2004). In mammalian cells, PTEN was also found to localise to the cytosol, membranes, nuclei, mitochondria and ER (Bononi et al., 2013; Bononi and Pinton, 2015; Putz et al., 2012; Zhu et al., 2009), which was not observed for *E. festucae* TepA.

MssD in *E. festucae* localised to the PM in a punctate pattern, with the signal being weak at the hyphal tip and increasingly stronger in older hyphae. In contrast, in *N. crassa*, Mss-4 localised to the membrane only subapical to the hyphal tip and at septa, as well as to intracellular filamentous structures (Mähs et al., 2012). In *A. thaliana* and tobacco root hairs and pollen tubes, PI4K5 localised to a sub-apical ring in actively growing cells and to the apex in non-growing cells (Ischebeck et al., 2008; Stenzel et al., 2008; Stenzel et al., 2011). However, *S. cerevisiae* Mss4 was found to localise evenly at the PM, indicating that localisation of Mss4 may not be universally conserved (Homma et al., 1998, Audhya and Emr, 2003; Vernay et al., 2012). Interestingly, the localisation of eGFP-MssD correlated with the localisation of the PI[4,5]P_2_ molecular probe in mature hyphae of *E. festucae*, but not at the hyphal tip. This could indicate that only low concentrations of MssD at the hyphal tip are required to generate the asymmetric distribution of PI[4,5]P_2_, or that a stable accumulation of this protein could not be observed due to the active nature of the endocytic collar.

Attempts to delete *E. festucae mssD* were unsuccessful, indicating that it is essential for fungal growth, as has been observed in *N. crassa* and in yeast (Desrivières et al., 1998; Mähs et al., 2012). Conditional mutants have demonstrated essential roles for Mss-4 in actin organization, hyphal morphogenesis and yeast to filamentous growth transition in *C. albicans* (Homma et al., 1998; Mähs et al., 2012; Vernay et al., 2012). Overexpression of *mssD* in *E. festucae* did not change the hyphal morphology in axenic culture, but the highest expression level obtained was approx. 6-fold relative to the WT *mssD* expression, which is low compared to the level of overexpression achieved with the same promoter for *tepA* (approx. 200 fold) or other proteins (Hassing et al., 2020). Thus, it is possible that overexpression of *mssD* negatively impacts on fungal growth.

Both *tepA* deletion and overexpression strains were found to be indistinguishable from WT in their hyphal growth rate and colony morphology, a result similar to what has been observed for deletion of this gene in *F. graminearum* and *U. maydis* (Vijayakrishnapillai et al., 2018; Zhang et al., 2010). TEM analysis revealed that the *S. pombe tep1* deletion mutant has an abnormal vacuolar morphology, highlighting the subtle nature of some culture phenotypes (Mitra et al., 2004).

Inoculation of the *E. festucae tepA* deletion mutants into *L. perenne* did not significantly change growth of the host plant when compared to a WT interaction. Similarly, *F. oxysporum ΔtepA* strains were equally virulent on tomato plants as WT. Interestingly, the corresponding TepA homologues in *F. graminearum* and *U. maydis* have been shown to be required for virulence on wheat and maize, respectively (Vijayakrishnapillai et al., 2018; Zhang et al., 2010). These species differences might be the result of slightly different infection processes, such as a requirement for chemotaxis.

Interestingly, leaves of *L. perenne* plants infected by two *mssD* overexpression strains were significantly longer than those of WT-infected plants which correlated with reduction in fungal biomass in plants infected with these two strains. A similar phenotype has previously been described for the *E. festucae cdc42* deletion mutant, where the increased length and reduced fungal biomass was due to defects in intercalary growth (Kayano et al., 2018). Whether the phenotype observed here for plants infected with *mssD* overexpression strains is also correlated with defects in intercalary growth remains to be elucidated.

Overexpression of *mssD* and deletion or overexpression of *tepA* did not appear to affect the localisation and concentration of the molecular probe for PI[4,5]P_2_. The absence of a detectable change in PI[4,5]P_2_ levels in the *E. festucae mssD* OE strains, in contrast to *MSS4* overexpression in *S. cerevisiae* (Desrivières et al., 1998), may have been due to the low overexpression levels. Given that the concentration of PI[3,4,5]P_3_ appears to be extremely low, it was perhaps not surprising that deletion or OE of *tepA* had no effect on the PI[4,5]P_2_ pool.

While the molecular probe for the detection of PI[3,4,5]P_3_ localised to the cytosol in *mssD* and *tepA* OE strains, in the *tepA* deletion strains it localised to septa, demonstrating for the first time the presence of this lipid species in filamentous fungi. Consistent with this result is the observation that TepA tagged with mCherry also localises to septa. This result demonstrates that PI[3,4,5]P_3_ is present albeit in very low quantities, in the cell during normal fungal growth. Similarly, in *S. pombe*, PI[3,4,5]P_3_ was only detected upon deletion of *tep1* (Balla, 2013; Mitra et al., 2004). As fungi lack homologues of the 3-kinase responsible for the phosphorylation of PI[4,5]P_2_, a pathway for the generation of PI[3,4,5]P_3_ similar to that described in *S. pombe* seems likely, where the production of this lipid species relies on the class III phosphatidylinositol 3-kinase, Vps34p, and the phosphatidylinositol 4-phosphate 5-kinase, Its3p (Balla, 2013; Mitra et al., 2004).

In the slime mould *Dictyostelium*, cells display a steep PI[3,4,5]P_3_ gradient, with highest concentrations of PI[3,4,5]P_3_ at the leading edge. Deletion of either the phosphoinositide 3-kinases or PTEN dissipated the PI[3,4,5]P_3_ gradient and reduced the efficiency of chemotaxis (Iijima and Devreotes, 2002; Iijima et al., 2004). Chemotaxis in mammalian neutrophils and macrophages also requires a PI[3,4,5]P_3_ gradient, although it appears to be regulated in a PTEN-independent way (Weiger and Parent, 2012). Here we found that chemotropic growth of *F. oxysporum* towards tomato root exudate or the chemoattractant horseradish peroxidase was impaired in *tepA* deletion mutants, uncovering for the first time a role for PI[3,4,5]P_3_ signalling in the control of directional hyphal growth. Moreover, our findings suggest that the role of PI[3,4,5]P_3_ in chemosensing is conserved across kingdoms. However, the exact functional link between PI[3,4,5]P_3_ and the downstream signalling components regulating chemotropic growth in *F. oxysporum*, such as the CWI MAPK cascade (Turrà et al., 2015), remains to be established. Likewise, it is currently unknown whether an asymmetric distribution of PI[3,4,5]P_3_ could govern chemotropism in this fungal pathogen, as described for *Dictyostelium* (Iijima and Devreotes, 2002; Iijima et al., 2004).

In summary, by analysing the localisation of PI species in the fungal endophyte *E. festucae*, we observed distinct localisation patterns for PI[3]P, PI[4]P, PI[3,5]P_2_ and PI[4,5]P_2_, and identified PI[3,4,5]P_3_, a PI species not previously reported in filamentous fungi. While deletion of the homologues of PTEN in *E. festucae* and *F. oxysporum* had no impact on the ability of these fungi to form mutualistic or pathogenic interactions with the plant hosts, we show that PTEN homologues are required for fungal chemotropism, potentially through establishment of a cellular PI[3,4,5]P_3_ gradient.

## Acknowledgements

This research was supported by grants from the Tertiary Education Commission to the Bio-Protection Research Centre, the Royal Society of New Zealand Marsden Fund (MAU1301) and by Massey University. Work in the Di Pietro lab was supported by by grants BIO2016-78923-R and PID2019-108045RB-I00 from the Spanish Ministerio de Ciencia e Innovación (MICINN). The authors thank Niki Murray, Pani Vijayan and Matthew Savoian (Manawatu Microscopy and Imaging Centre) for support with microscopy and imaging, Arvina Ram for technical support, David Winter for advice on data analysis, Pierre Dupont for bioinformatics assistance, Kimberly Green for help with some of the microscopy, and Chris Schardl for providing *Epichloë* spp. sequence data.

## Author Contributions

BH, CJE, CHM, ADP and BS planned and designed the research. BH, AC, TRF and CJE performed the experiments. BH, AC, TRF, CJE, CHM, ADP and BS analyzed the results. CJE, CHM, ADP and BS supervised the project. BH, TRF, CHM, ADP and BS wrote the manuscript.

