## Supplementary material for "A homologue of the mammalian tumour suppressor protein PTEN is a functional lipid phosphatase and required for chemotaxis in filamentous fungi": Supplmentary Data

New Phytologist

***New Phytologist* Supporting Information**

Article acceptance date: Click here to enter a date.

The following Supporting Information is available for this article:

**Fig. S1: Verification of the expression of molecular probes in *Epichloë festucae* by western blot analysis.**

**Fig. S2: Localisation of the PI[3,4]$P_2$ and PI[3,4,5]$P_3$ molecular probes in *Epichloë festucae* grown in axenic culture.**

**Fig. S3: Asymmetric distribution of the PI[4,5]$P_2$ molecular probe at the *Epichloë festucae* hyphal tip.**

**Fig. S4: Verification of the expression of eGFP-MssD, eGFP-TepA and TepA-mCherry fusion proteins in *Epichloë festucae* by western blot analysis.**

**Fig. S5: Targeted replacement of the *Epichloë festucae tepA* gene.**

**Fig. S6: Expression analysis of *Epichloë festucae mssD* and *tepA* overexpression strains by qRT-PCR.**

**Fig. S7: Culture phenotype of *Epichloë festucae* wild-type, *mssD* overexpression and *tepA* deletion and overexpression mutants.**

**Fig. S8: Growth analysis of *Lolium perenne* plants infected with *Epichloë festucae* wild-type and *tepA* overexpression strains.**

**Fig. S9: Analysis of the *in planta* fungal biomass of *Epichloë festucae* wild-type and *mssD* overexpression strains.**

**Fig. S10: Confocal depth series of longitudinal sections of *Lolium perenne* infected with *Epichloë festucae* wild-type and *mssD* and *tepA* mutant strains.**

**Fig. S11: Targeted replacement of the *Fusarium oxysporum tepA* gene.**

**Fig. S12: TepA is not required for virulence of *Fusarium oxysporum*.**

**Table S1: Biological material.**

New Phytologist

**Table S2: Primers used in this study.**

**Methods S1: Construct design.**

a

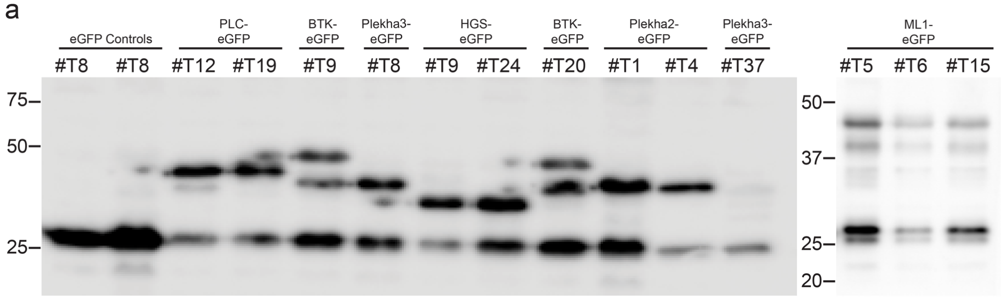

b

| Construct | Domain Type | Phospholipid | Strain | Predicted MW (KDa) |
|---|---|---|---|---|
| eGFP | Control | | 8 | 29.95 |
| PLC-eGFP | PH | $PI[4,5]P_2$ | 12 | 42.34 |
| | | | 19 | |
| BTK-eGFP | PH | $PI[3,4,5]P_3$ | 9 | 47.13 |
| Plekha3-eGFP | PH | PI[4]P | 8 | 38.19 |
| HGS-eGFP | FYVE | PI[3]P | 9 | 32.26 |
| | | | 24 | |
| BTK-eGFP | PH | $PI[3,4,5]P_3$ | 20 | 47.13 |
| Plekha2-eGFP | PH | $PI[3,4]P_2$ | 1 | 40.01 |
| | | | 4 | |
| Plekha3-eGFP | PH | PI[4]P | 37 | 38.19 |
| ML1-eGFP | N-terminal lipid binding domain | $PI[3,5]P_2$ | 5 | 43 |
| | | | 6 | |
| | | | 12 | |

c

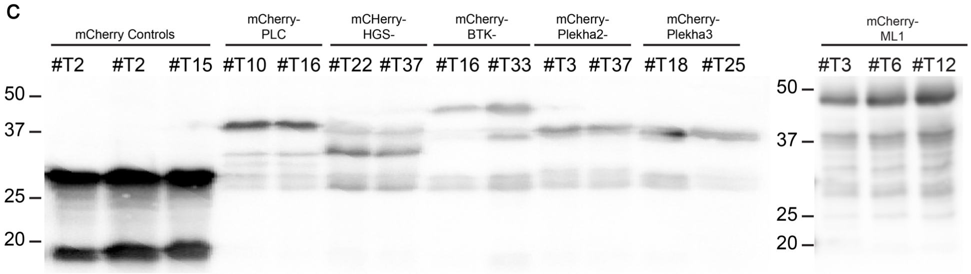

d

| Construct | Domain Type | Phospholipid | Strain | Predicted MW (KDa) |
|---|---|---|---|---|
| mCherry | Control | | 2 | 26.73 |
| | | | 15 | |
| mCherry-PLC | PH | $PI[4,5]P_2$ | 10 | 42.04 |
| | | | 16 | |
| mCherry-HGS | FYVE | PI[3]P | 22 | 34.97 |
| | | | 37 | |
| mCherry-BTK | PH | $PI[3,4,5]P_3$ | 16 | 46.84 |
| | | | 33 | |
| mCherry-Plekha2 | PH | $PI[3,4]P_2$ | 3 | 39.72 |
| | | | 37 | |
| mCherry-Plekha3 | PH | PI[4]P | 18 | 38.03 |
| | | | 25 | |
| mCherry-ML1 | N-terminal lipid binding domain | $PI[3,5]P_2$ | 4 | 42.78 |
| | | | 6 | |
| | | | 16 | |

**Fig. S1: Verification of the expression of molecular probes in *Epichloë festucae* by western blot analysis.**

(a) Western blot of total protein extract of lipid-binding molecular probe-expressing strains probed with the anti-GFP antibody (Abcam). Total protein extract of strain PN4175 expressing cytosolic eGFP (pCT74) was used as a control. For each strain, 50 µg of total protein was loaded and samples separated on a 10% SDS PAGE gel. (b) Table of molecular probe constructs and their expected molecular weight (kDa) as calculated using https://www.bioinformatics.org/sms/prot_mw.html. (c) Western blot of total protein extract of lipid-binding molecular probe-expressing strains probed with the anti-mCherry antibody (Abcam). For each strain, 50 µg of total protein was loaded and samples were separated on a 10% SDS PAGE gel. (d) Table of molecular probe constructs and their expected molecular weight (kDa) as calculated using https://www.bioinformatics.org/sms/prot_mw.html.

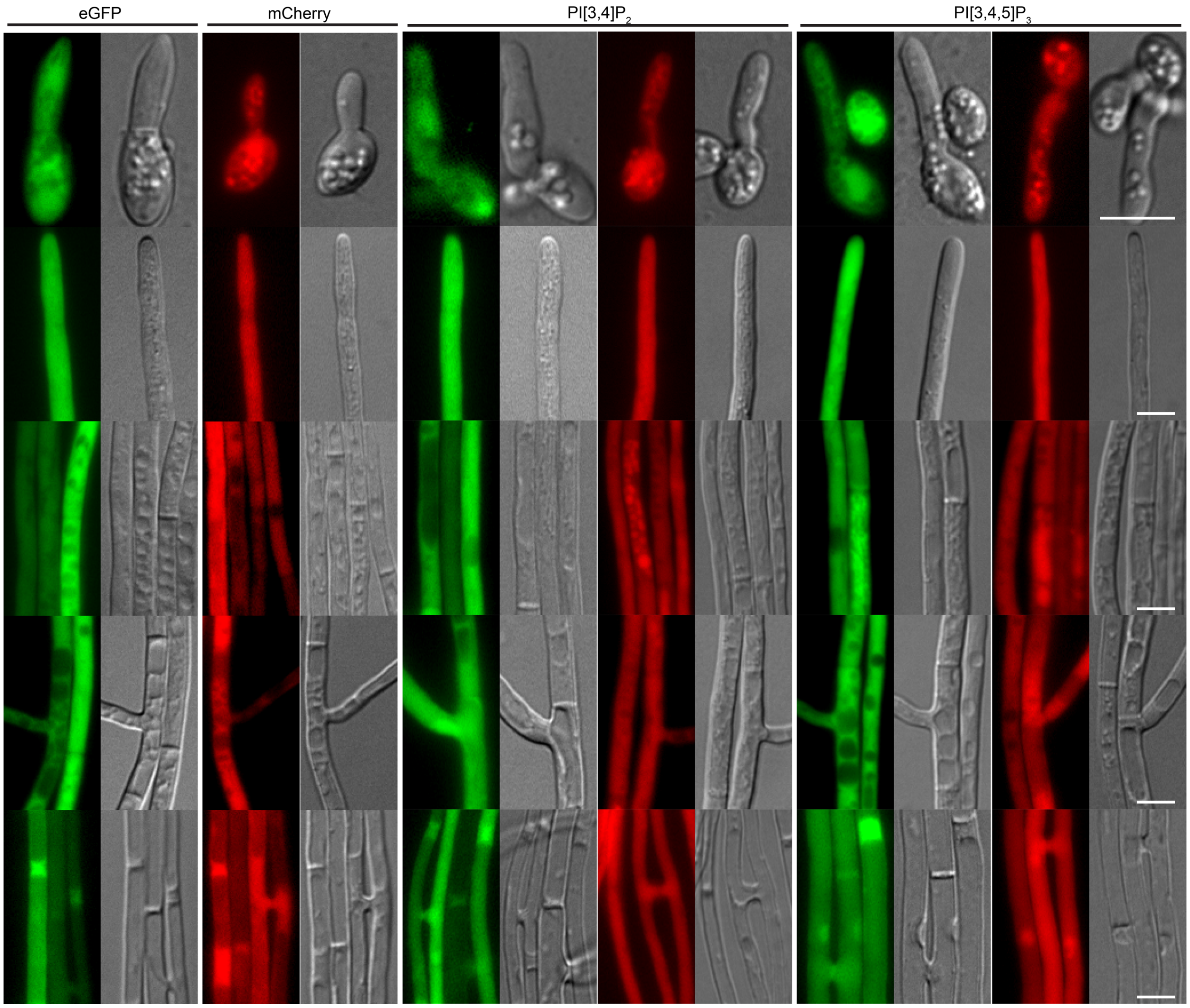

**Fig. S2: Localisation of the PI[3,4]$P_2$ and PI[3,4,5]$P_3$ molecular probes in *Epichloë festucae* grown in axenic culture.**

Strains were grown on 1.5% $H_2O$ agar for 5 d before examination using a fluorescence microscope. Images shown are representative of all strains analysed and show localization of the molecular probes in hyphae of different ages. Images of spores were acquired from *E. festucae* E2368 transformants, whereas all others are from Fl1. Cytosolic eGFP (pCE25): #T5 (E2368), #T8 (Fl1); cytosolic mCherry (pCE126): #T7 (E2368), #T2 (Fl1); Plekha2-eGFP (PI[3,4]$P_2$ (pCE108): #T7 (E2368), #T1 (Fl1); mCherry-Plekha2 (PI[3,4]$P_2$, pCE113): #T6 (E2368), #T37 (Fl1); BTK-eGFP (PI[3,4,5]$P_3$, pCE107): #T14 (E2368), #T9 (Fl1); mCherry-BTK (PI[3,4,5]$P_3$, pCE112): #T1 (E2368), #T16 (Fl1); Bar = 5 µm.

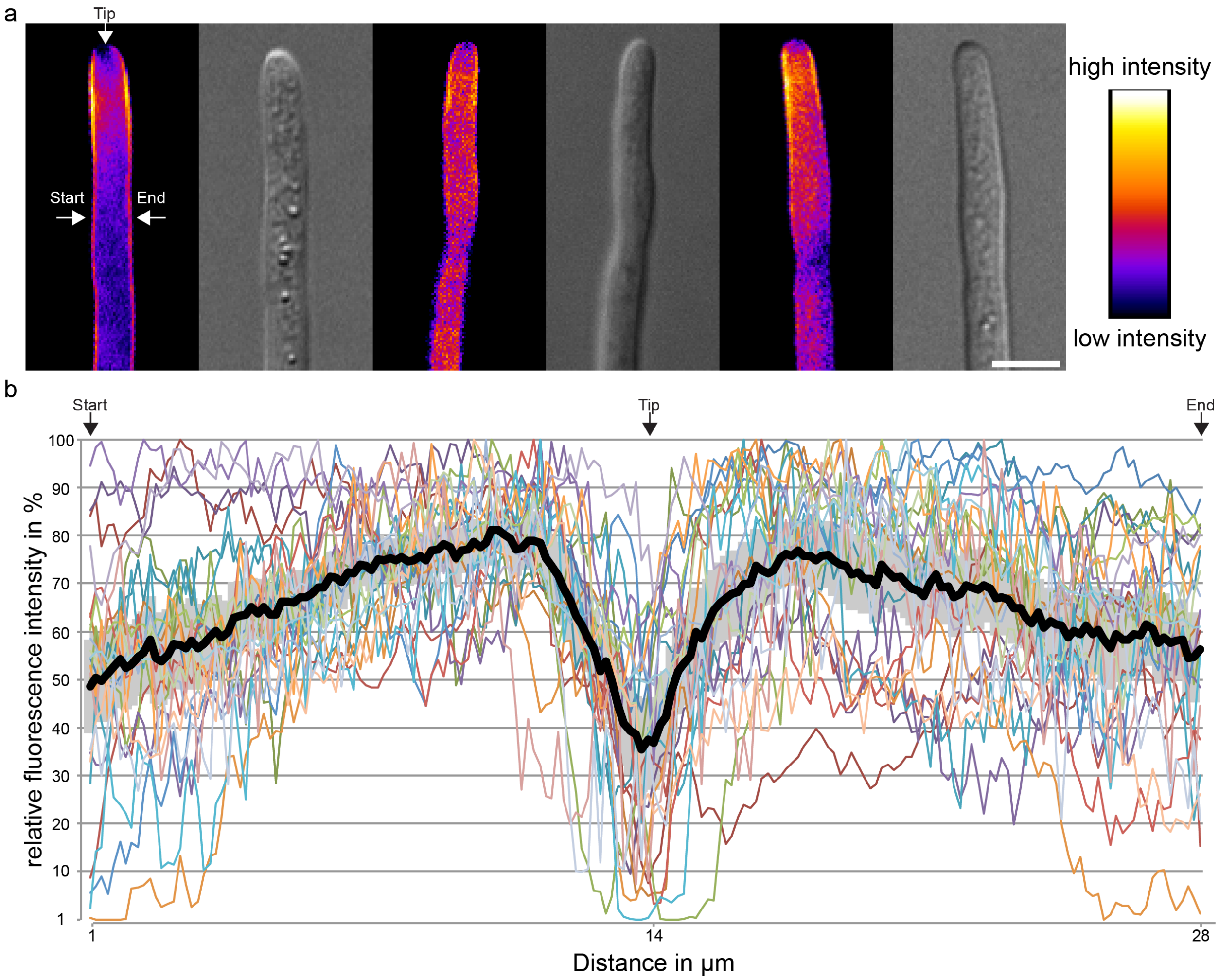

**Fig. S3: Asymmetric distribution of the PI[4,5]$P_2$ molecular probe at the *Epichloë festucae* hyphal tip.**

(a) Representative differential interference contrast and fluorescent microscopy images of the PI[4,5]$P_2$ biosensor at hyphal tips, pseudocoloured to indicate the saturation of each pixel from 0 to 255 (fully saturated). A three-pixel-wide line was drawn along the cell membrane from the start arrow around the tip to the end arrow. Graphs represent the concentration of biosensor along this line. Cultures were grown on 1.5% (w/v) water agar for 6-9 d. Bar = 5 µm. (b) Scatter graph summarising the concentration of the PI[4,5]$P_2$ biosensor along the cell membrane of 24 hyphal tips. Black line = trend line.

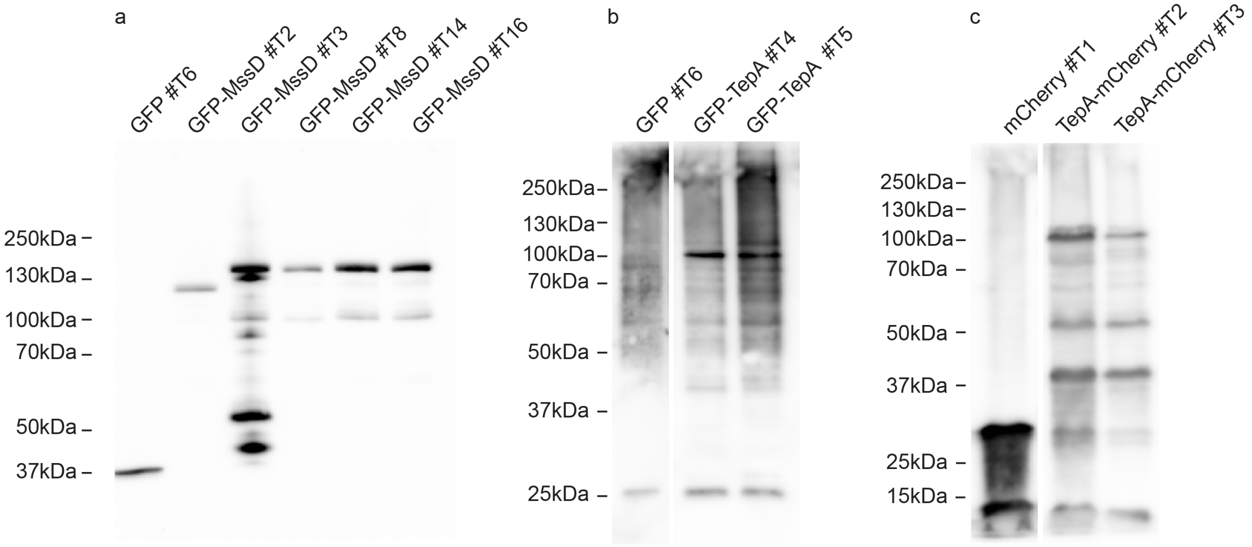

**Fig. S4: Verification of the expression of eGFP-MssD, eGFP-TepA and TepA-mCherry fusion proteins in *Epichloë festucae* by western blot analysis.**

Strains expressing the fusion proteins were incubated in liquid media for 3 d and total protein extracted. Approx. 15 µl of protein extract was loaded onto a 10% SDS acrylamide gel, separated and transferred onto PVDF membrane. Following transfer, membranes were probed with anti-GFP or anti-mCherry antibodies (Abcam), respectively. Following the incubation with a secondary anti-rabbit antibody (Abcam) conjugated to horseradish peroxidase (HRP), western blots were developed using a chemiluminescence reaction. All controls and fusion-protein constructs were expressed in the WT background, and the detection of eGFP (GFP#T6), eGFP-fusion proteins (eGFP-TepA#T4 and eGFP-TepA#T5), mCherry (mCherry#T1) and mCherry-fusion proteins (TepA-mCherry#T2 and #T3) is shown. The eGFP and mCherry expression controls are the same as previously described in Hassing *et al*. 2019 as they were run on the same gel. Expected protein sizes: GFP: 27 kDa; mCherry: 28.8 kDa; GFP-TepA: 86.18 kDa; TepA-mCherry: 87.98 kDa.

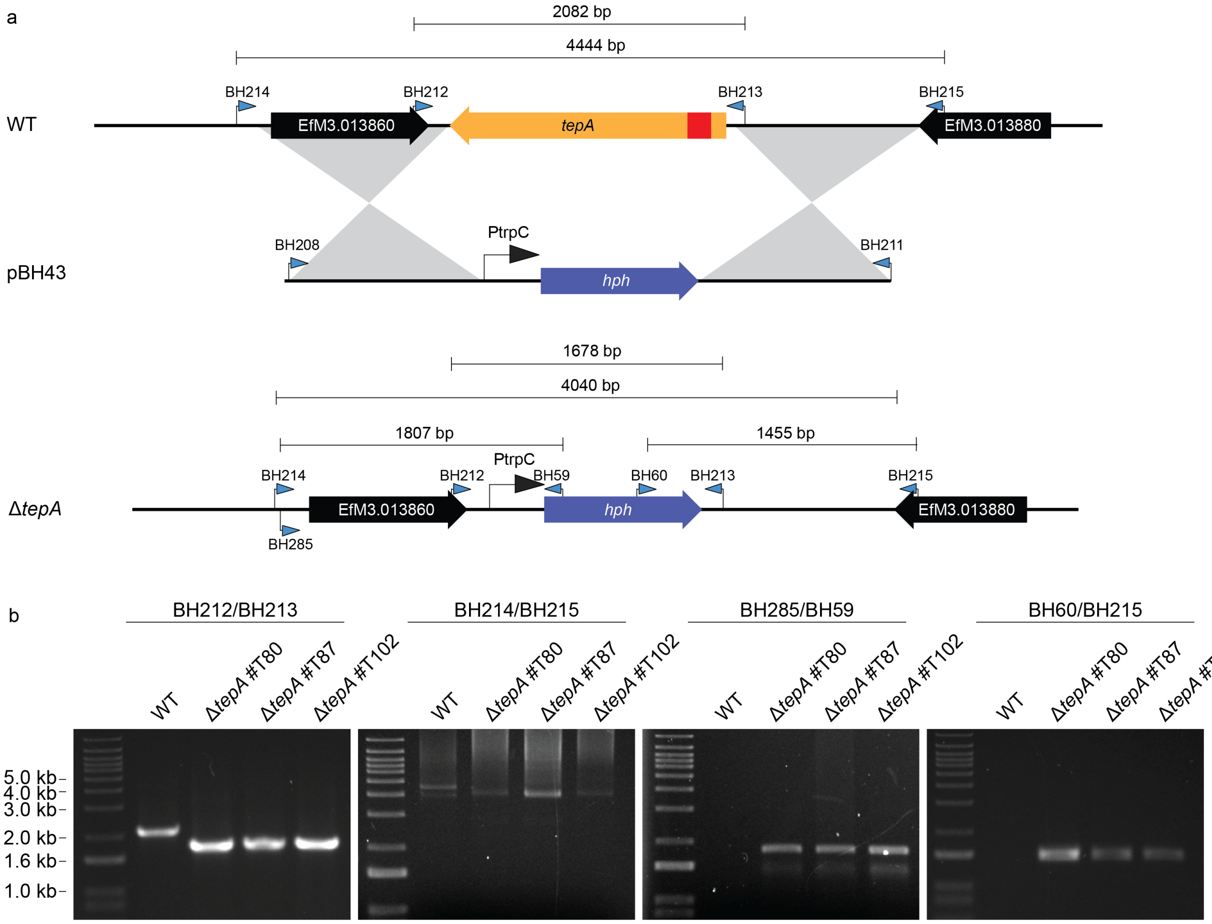

**Fig. S5: Targeted replacement of the *Epichloë festucae tepA* gene.**

(a) Physical map of the *tepA* wild-type (WT) genomic locus, linear insert of the *tepA* replacement construct, pBH43, and the mutant locus. Grey shading indicates regions of recombination. Numbers indicate the PCR primer pairs used for Gibson assembly (BH208/BH211) and deletion mutant screening (BH59/BH60/ BH212/BH213/BH214/BH215/BH285). (b) PCR screening of deletion candidates with the PCR primer pair BH212/BH213, generated expected bands of approx. 1.7 kb for the deletion mutants and approx. 2.1 kb for the WT strain. The primer pair BH214/BH215 generated expected bands of approx. 4.0 kb for the deletion mutants, and 4.4 kb for the WT strains. The primer pair BH60/BH215 generated a band of 1.5 kb and the primer pair BH285/BH59 a band of approx. 1.8 kb for the deletion mutants.

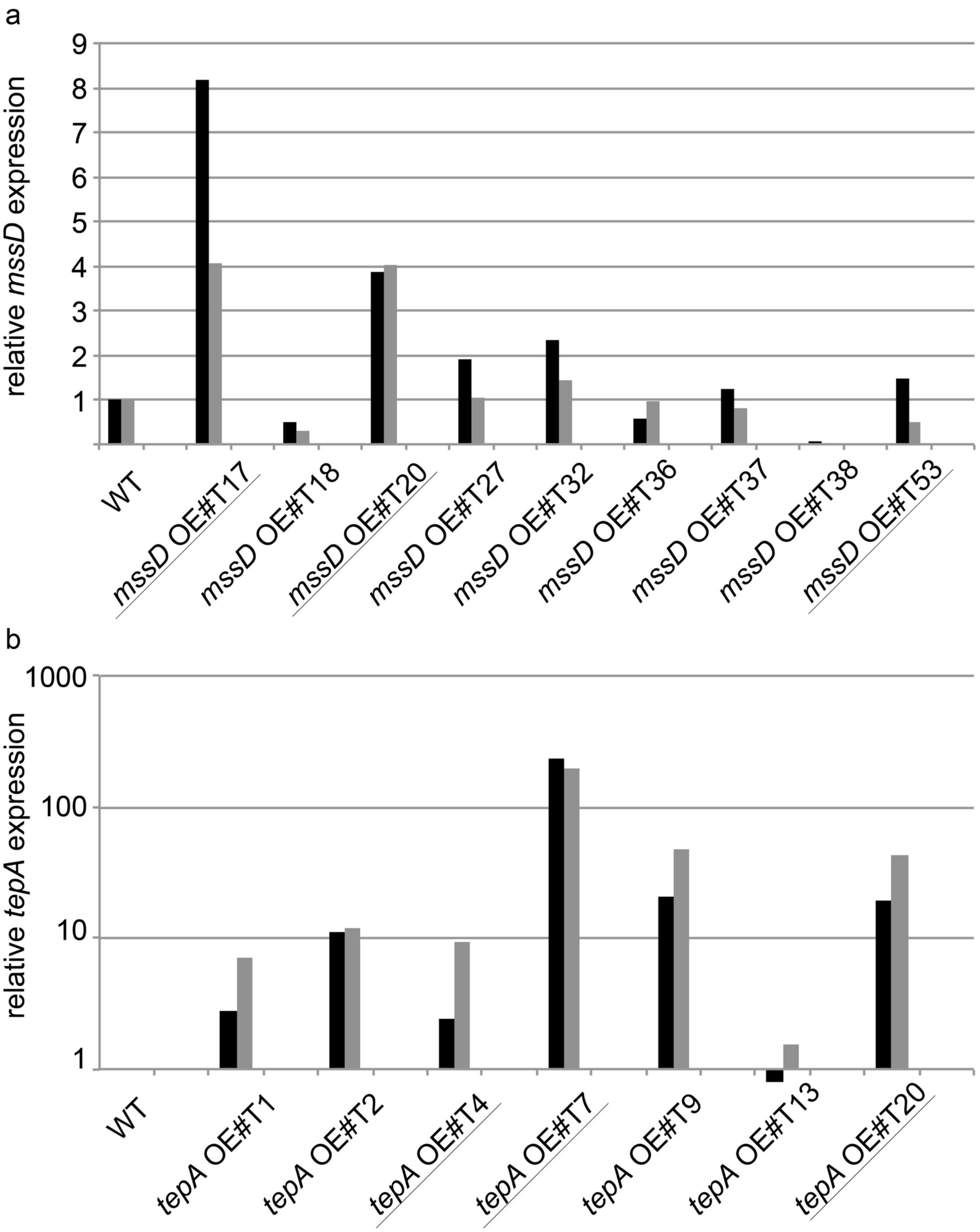

**Fig. S6: Expression analysis of *Epichloë festucae mssD* and *tepA* overexpression strains by RT-qPCR.**

Expression of *mssD* (a) and *tepA* (b) was determined relative to the wild-type (WT) expression using genes coding for elongation factor 2 (*EF-2*, black bar) and 40S ribosomal protein S22 (*S22*, grey bar) for normalisation with two technical replicates. For the amplification of *mssD*, the primer pair AC33/43, and for *tepA*, the primer pair BH189/199, was used. The relative expression was calculated as previously described (Lukito et al., 2015). Overexpression (OE) strains chosen for further analysis are underlined. Note the different scales on the y-axis.

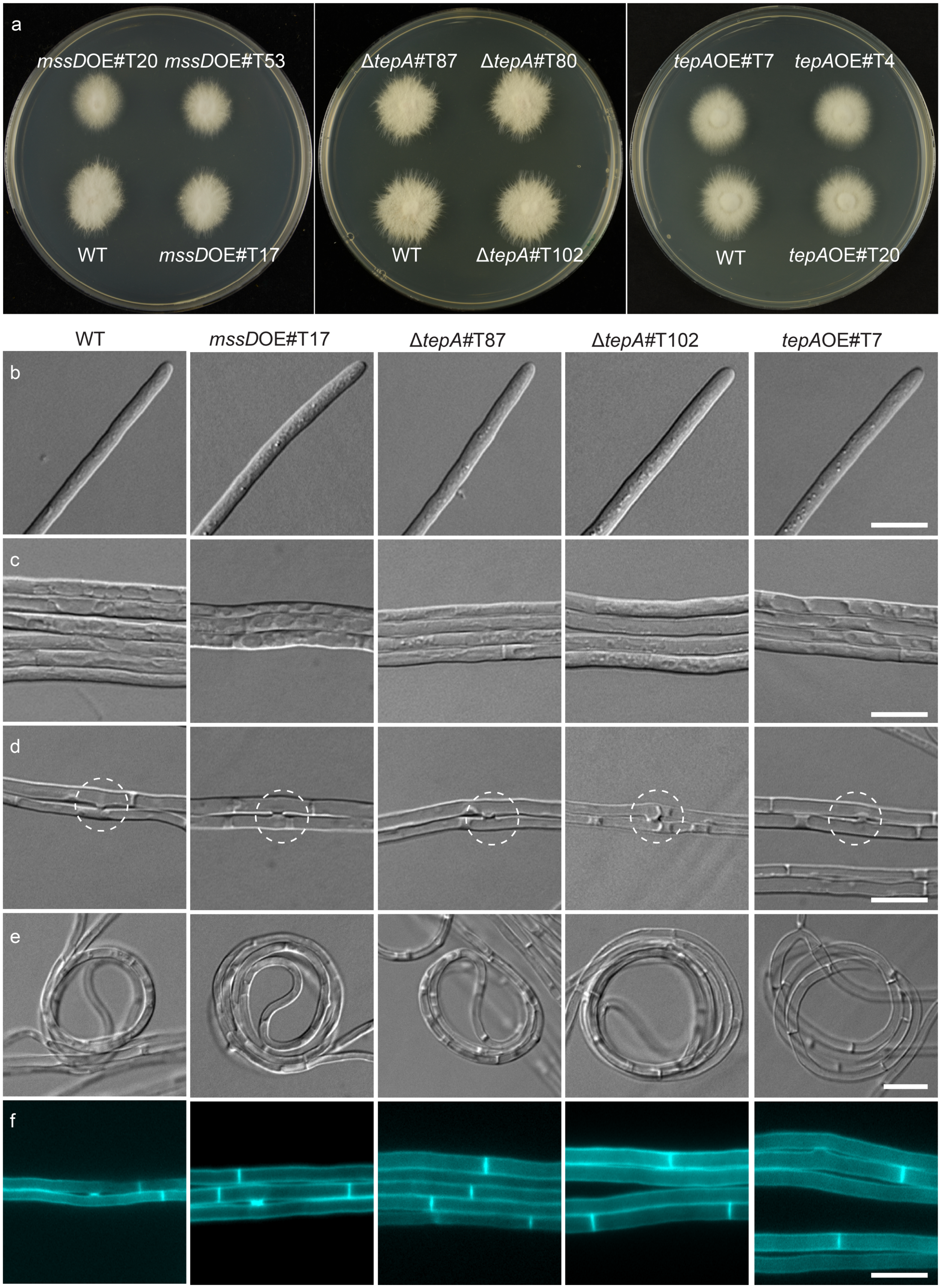
a
mssDOE#T20 mssDOE#T53
WT mssDOE#T17
ΔtepA#T87 ΔtepA#T80
WT ΔtepA#T102
tepAOE#T7 tepAOE#T4
WT tepAOE#T20
WT
mssDOE#T17
ΔtepA#T87
ΔtepA#T102
tepAOE#T7
b
c
d
e
f

**Fig. S7: Culture phenotype of *Epichloë festucae* wild-type, *mssD* overexpression and *tepA* deletion and overexpression mutants.**

Wild-type (WT), *mssD* overexpression strains (#T17, #T20, #T53), *tepA* deletion strains (#T80, #T87, #T102) and *tepA* overexpression (OE) strains (#T4, #T7, #T20) were analysed multiple times. The images shown were captured of *mssD* OE #T17, Δ*tepA* #T87, Δ*tepA* #T102 and *tepA* OE #T7 and are representative of all strains analysed. (a) Whole colony morphology of WT and mutant strains grown on 2.4% PD agar for 7 d before examination; (b-f) strains were grown for 7 d on 1.5% $H_2O$ agar before analysis; (b) Hyphal tip morphology in WT and mutant strains; (c) Formation of hyphal bundles in WT and mutant strains; (d) Hyphal fusion in WT and mutant strains; (e) Hyphal coil formation in WT and mutant strains; (f) Typical staining of hyphal bundle with Calcofluor white (CFW). White circle: hyphal fusion; Bar = 10 µm.

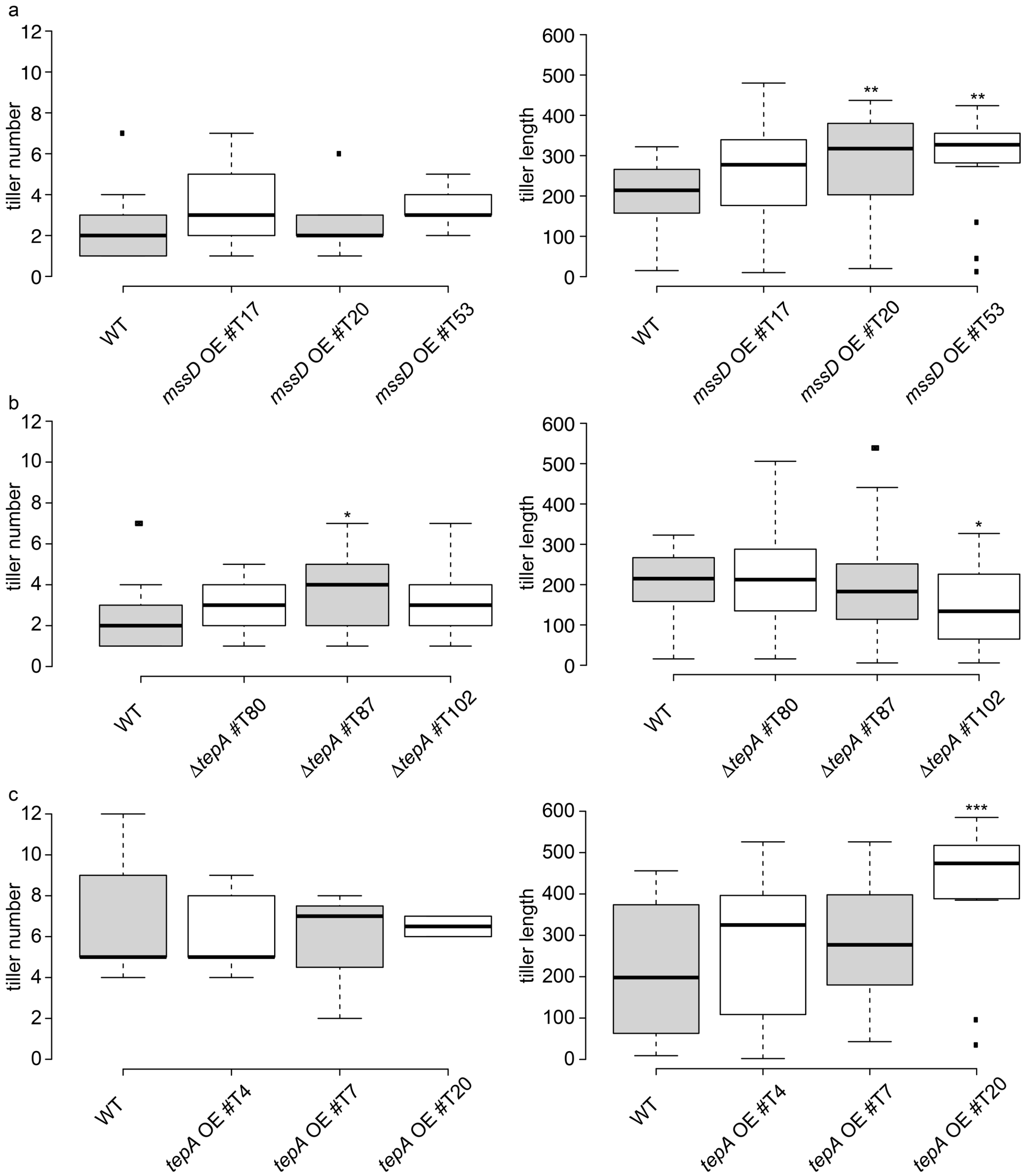

**Fig. S8: Growth analysis of *Lolium perenne* plants infected with *Epichloë festucae* wild-type and *tepA* overexpression strains.**

Tiller number of infected plants: wild-type (WT) (23), *mssD* overexpression strains #T17, #T20, #T53 (n=18/15/6), *tepA* deletion strains #T80, #T87, #T102 (n=31/32/18), WT (5), *tepA* overexpression strains #T4, #T7, #T20: (n=5/3/2). Box plots were generated using BoxPlotR (available on http://shiny.chemgrid.org/boxplotr/). One-way ANOVAs were used to test for differences in plant phenotypes between WT and mutant strains. In each case, the ANOVA was fitted with R and a Bonferroni correction was applied to all *p*-values to account for multiple testing: *$p \leq 0.05$; **$p \leq 0.01$; ***$p \leq 0.001$. All other differences are not significant. The highly significant difference regarding the tiller length of the *tepA* OE strain #T20 has not been mentioned in the main text due to the low and therefore statistically insignificant number of biological replicates.

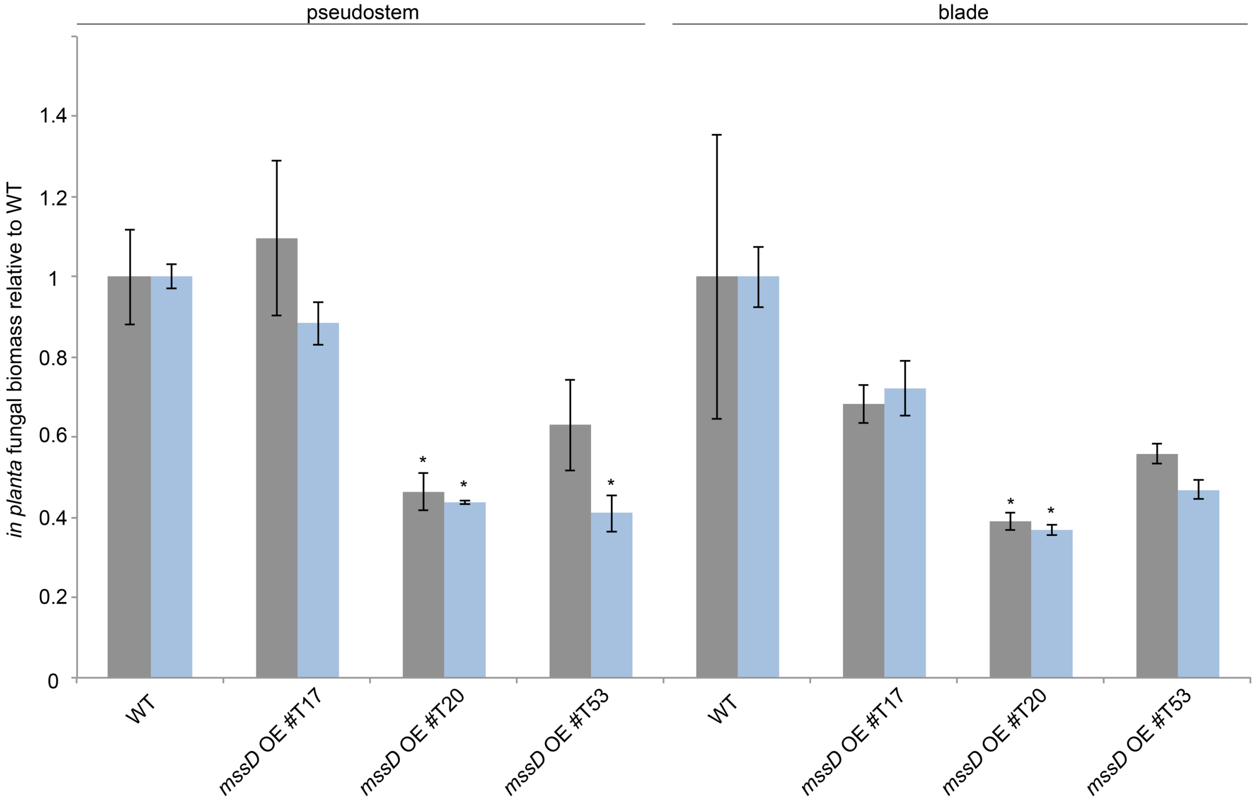

**Fig. S9: Analysis of the *in planta* fungal biomass of *Epichloë festucae* wild-type and *mssD* overexpression strains.**

The fungal biomass was analysed in three biological replicates of mature, infected *L. perenne* plants and is depicted relative to the WT biomass in the corresponding dataset. Grey bars show the relative biomass based on the presence of the fungal gene *hepA* (YL120F/YL120R) relative to the *L. perenne* gene *LpCCR1* (YL501F/YL501R) and blue bars the relative biomass based on the presence of the fungal gene *pacC* (YL113F/YL113R) relative to the *L. perenne* gene *LpCCR1* (YL502F/YL502R). Error bars represent the standard error. Asterisks represent statistically significant differences in the mutant-infected plants ($p \leq 0.05$) compared to the corresponding WT-infected plants calculated by a two-tailed Student *t*-Test. While the biomass was always significantly lower in *mssD* OE #T20-infected plants compared to WT-infected plants, the fungal biomass was only occasionally significantly reduced in *mssD* OE #T53-infected plants.

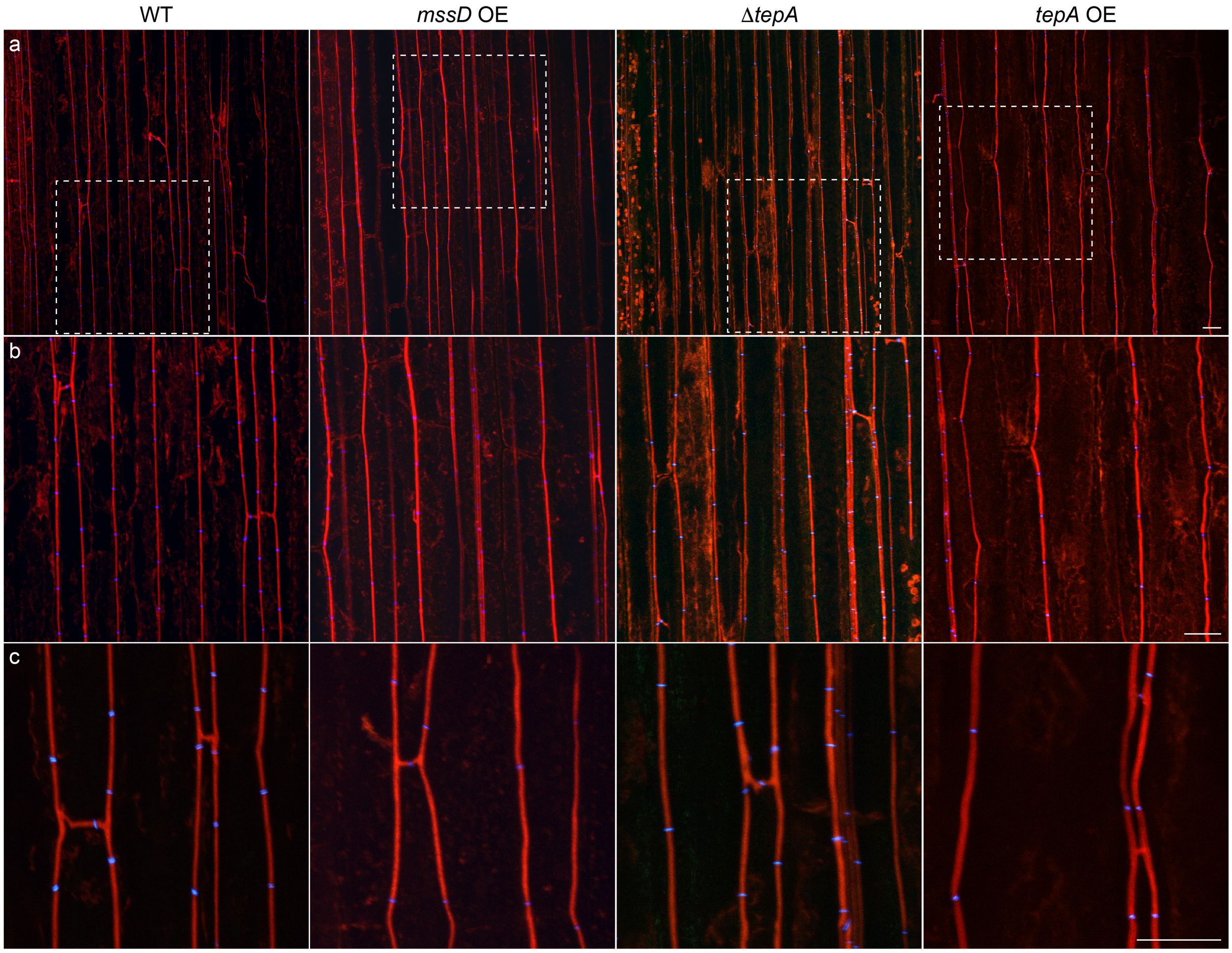

**Fig. S10: Confocal depth series of longitudinal sections of *Lolium perenne* infected with *Epichloë festucae* wild-type and *mssD* and *tepA* mutant strains.**

Infected *L. perenne* pseudostem samples were labelled with WGA-AF488 (chitin-binding, blue pseudocolour) and aniline blue (β-glucan-binding, red pseudocolour) and visualized with confocal scanning laser microscopy. Representative images of plant material infected with wild-type (WT), *mssD* overexpression (OE) #T17, Δ*tepA* #T87 and *tepA* OE #T7 strains. (a) Representative image (z = 6 µm) of strains *in planta;* Bar = 10 µm; (b) Magnified images of white boxed region in (a); Bar = 10 µm. (c) Higher magnification images (z = 4 µm) of hyphal branching and fusion. Bar = 10 µm.

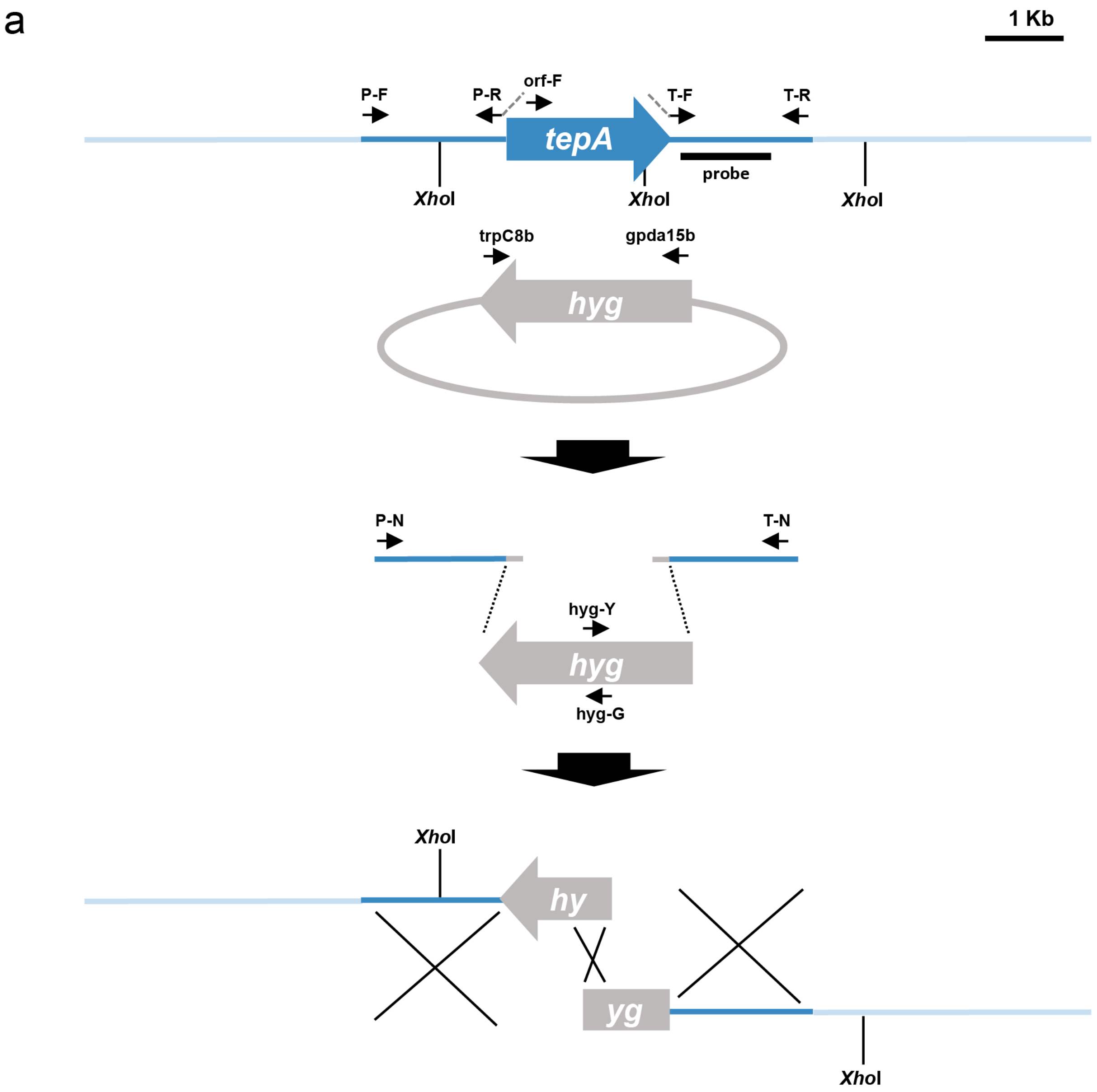

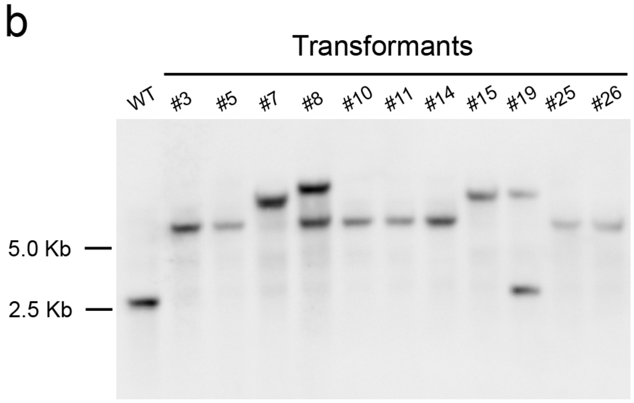

**Fig. S11: Targeted replacement of the *Fusarium oxysporum tepA* gene.**

(a) Physical maps of the *F. oxysporum tepA* locus and the split-marker gene replacement constructs obtained by fusion PCR. Relative positions of restriction sites and PCR primers are indicated. *hyg*, hygromycin resistance gene. (b) Southern blot analysis. Genomic DNA of the wild-type strain and eleven independent transformants was treated with *Xho*I, separated on a 0.7% agarose gel, transferred to a nylon membrane and hybridized with the DNA probe indicated in (a).

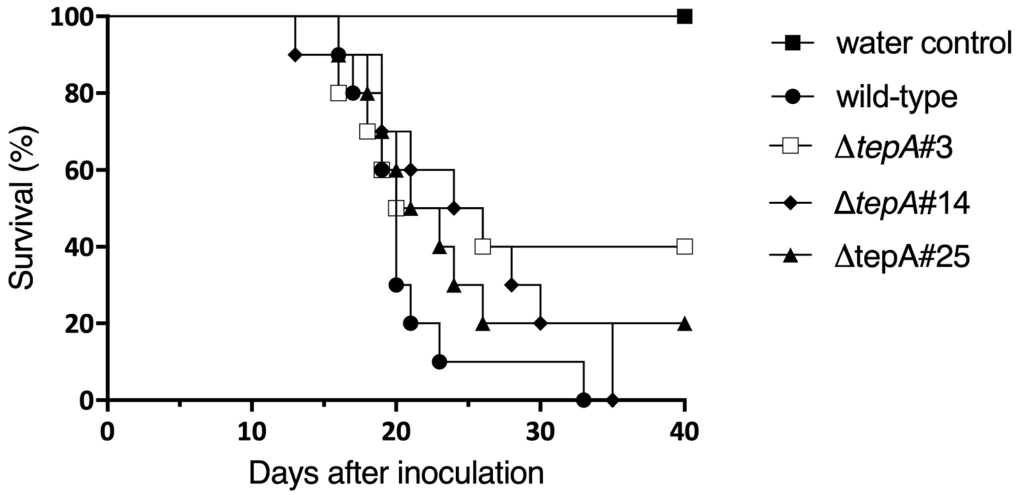

**Fig. S12: TepA is not required for virulence of *Fusarium oxysporum*.**
Roots of 2-week-old tomato plants were inoculated with 5 x $10^6$ microconidia $ml^{-1}$ of the indicated *F. oxysporum* strains and planted in minipots. Plant survival was recorded daily for 40 d. Data shown are from one representative experiment. All experiments were performed twice with similar results.

### Table S1: Biological material.

| Organism/Strain | Characteristics | Reference |
|---|---|---|
| ***E. coli*** | | |
| DH5α | F$^-$, ϕ80*lacZ*, ΔM15, Δ(*lacZYA-argF*), U169, *recA1*, *endA1*, *hsdR17* ($r_K^-$, $m_K^-$), *phoA*, *supE44*, λ$^-$, *thi-1*, *gyrA96*, *relA1* | Invitrogen |
| ***E. festucae*** | | |
| PN2278 (Fl1) | Wild-type isolated from *Festuca longifolia* | (Young et al., 2005) |
| E2368 | | (Schardl et al., 2013) |
| Δ*tepA* #T80 | WT/Δ*tepA::*P*trpC-hph;* Hyg$^R$ | This study |
| PN3278 (Δ*tepA* #T87) | WT/Δ*tepA::*P*trpC-hph;* Hyg$^R$ | This study |
| PN3279 (Δ*tep*A #T102) | WT/Δ*epA::*P*trpC-hph;* Hyg$^R$ | This study |
| Δ*tepA* #T87 PLC-GFP #TN1 | Δ*tepA*#T87/pCE105/pII99, Hyg$^R$, Gen$^R$ | This study |
| Δ*tepA* #T87 PLC-GFP #TN2 | Δ*tepA*#T87/pCE105/pII99, Hyg$^R$, Gen$^R$ | This study |
| Δ*tepA* #T87 BTK-GFP #15 | Δ*tepA*#T87/pCE107/pII99, Hyg$^R$, Gen$^R$ | This study |
| Δ*tepA* #T102 BTK-GFP #15 | Δ*tepA*#T102/pCE107/pII99, Hyg$^R$, Gen$^R$ | This study |

| | | |
|---|---|---|
| *tepA* overexpression #T4 | WT/pCE122, pSF16.17, Gen$^R$ | This study |
| *tepA* overexpression #T7 | WT/pCE122, pSF16.17, Gen$^R$ | This study |
| *tepA* overexpression #T20 | WT/pCE122, pSF16.17, Gen$^R$ | This study |
| *tepA* overexpression #T7 PLC-GFP #T6 | *tepA* overexpression#T7/pCE105/pBH12, Hyg$^R$, Gen$^R$ | This study |
| *tepA* overexpression #T7 PLC-GFP #T11 | *tepA* overexpression#T7/pCE105/pBH12, Hyg$^R$, Gen$^R$ | This study |
| *tepA* overexpression #T7 BTK-GFP #T1 | *tepA* overexpression#T7/pCE107/pBH12, Hyg$^R$, Gen$^R$ | This study |
| *tepA* overexpression #T7 BTK-GFP #T2 | *tepA* overexpression#T7/pCE107/pBH12, Hyg$^R$, Gen$^R$ | This study |
| *mssD* overexpression #T17 | WT/pCE101, pSF16.17, Gen$^R$ | This study |
| *mssD* overexpression #T20 | WT/pCE101, pSF16.17, Gen$^R$ | This study |
| *mssD* overexpression #T53 | WT/pCE101, pSF16.17, Gen$^R$ | This study |
| *mssD* overexpression #T17 PLC-GFP #T20 | WT/pCE101, pSF16.17, pCE105, pBH12, Gen$^R$, Hyg$^R$ | This study |
| *mssD* overexpression #T17 BTK-GFP #T1 | WT/pCE101, pSF16.17, pCE105, Gen$^R$, Hyg$^R$ | This study |
| *mssD* overexpression #T17 BTK-GFP #T8 | WT/pCE101, pSF16.17, pCE105, Gen$^R$, Hyg$^R$ | This study |
| WT GFP-TepA #T4 | WT/pBH41, Gen$^R$ | This study |
| WT GFP-TepA #T5 | WT/pBH41, Gen$^R$ | This study |
| WT GFP-TepA #T9 | WT/pBH41, Gen$^R$ | This study |
| WT GFP-TepA #T15 | WT/pBH41, Gen$^R$ | This study |
| WT TepA-mCherry #T1 | WT/pBH42, Hyg$^R$ | This study |
| WT TepA-mCherry #T2 | WT/pBH42, Hyg$^R$ | This study |
| WT TepA-mCherry #T3 | WT/pBH42, Hyg$^R$ | This study |
| WT TepA-mCherry #T10 | WT/pBH42, Hyg$^R$ | This study |
| WT GFP-MssD #T2 | WT/pBH72, Hyg$^R$ | This study |
| WT GFP-MssD #T3 | WT/pBH72, Hyg$^R$ | This study |
| WT GFP-MssD #T8 | WT/pBH72, Hyg$^R$ | This study |
| WT GFP-MssD #T14 | WT/pBH72, Hyg$^R$ | This study |
| WT GFP-MssD #T16 | WT/pBH72, Hyg$^R$ | This study |
| WT GFP #T6 | WT/pBH28, Gen$^R$ | (Hassing et al., 2020) |
| WT GFP #T2 | WT/pCE125, pSF16.17, Gen$^R$ | This study |
| WT GFP #T8 | WT/pCE125, pSF16.17, Gen$^R$ | This study |
| WT mCherry #T1 | WT/pCE126, pSF16.17, Gen$^R$ | This study |
| WT mCherry #T2 | WT/pCE126, pSF16.17, Gen$^R$ | This study |
| WT PLC-GFP #T6 | WT/pCE105/pSF16.17, Gen$^R$ | This study |
| WT PLC-GFP #T12 | WT/pCE105/pSF16.17, Gen$^R$ | This study |
| WT PLC-GFP #T19 | WT/pCE105/pSF16.17, Gen$^R$ | This study |
| WT PLC-GFP #T2 | WT/pCE105/pBH12, Hyg$^R$ | This study |
| WT PLC-GFP #T3 | WT/pCE105/pBH12, Hyg$^R$ | This study |
| WT PLC-GFP #T7 | WT/pCE105/pBH12, Hyg$^R$ | This study |
| WT mCherry-PLC #T10 | WT/pCE110/pSF16.17, Gen$^R$ | This study |
| WT mCherry-PLC #T16 | WT/pCE110/pSF16.17, Gen$^R$ | This study |
| WT BTK-GFP #T9 | WT/pCE107/pSF16.17, Gen$^R$ | This study |
| WT BTK-GFP #T20 | WT/pCE107/pSF16.17, Gen$^R$ | This study |
| WT mCherry-BTK #T16 | WT/pCE112/pSF16.17, Gen$^R$ | This study |
| WT mCherry-BTK #T32 | WT/pCE112/pSF16.17, Gen$^R$ | This study |
| WT mCherry-BTK #T33 | WT/pCE112/pSF16.17, Gen$^R$ | This study |
| WT HGS-GFP #T9 | WT/pCE106/pSF16.17, Gen$^R$ | This study |
| WT HGS-GFP #T18 | WT/pCE106/pSF16.17, Gen$^R$ | This study |
| WT HGS-GFP #T24 | WT/pCE106/pSF16.17, Gen$^R$ | This study |

| | | |
|---|---|---|
| WT mCherry-HGS #T5 | WT/pCE111/pSF16.17, $Gen^R$ | This study |
| WT mCherry-HGS #T22 | WT/pCE111/pSF16.17, $Gen^R$ | This study |
| WT mCherry-HGS #T37 | WT/pCE111/pSF16.17, $Gen^R$ | This study |
| WT PLEKHA2-GFP #T1 | WT/pCE108/pSF16.17, $Gen^R$ | This study |
| WT PLEKHA2-GFP #T4 | WT/pCE108/pSF16.17, $Gen^R$ | This study |
| WT mCherry-PLEKHA2 #T3 | WT/pCE113/pSF16.17, $Gen^R$ | This study |
| WT mCherry-PLEKHA2 #T37 | WT/pCE113/pSF16.17, $Gen^R$ | This study |
| WT mCherry-PLEKHA2 #T40 | WT/pCE113/pSF16.17, $Gen^R$ | This study |
| WT PLEKHA3-GFP #T3 | WT/pCE109/pSF16.17, $Gen^R$ | This study |
| WT PLEKHA3-GFP #T8 | WT/pCE109/pSF16.17, $Gen^R$ | This study |
| WT PLEKHA3-GFP #T37 | WT/pCE109/pSF16.17, $Gen^R$ | This study |
| WT mCherry-PLEKHA3 #T18 | WT/pCE114/pSF16.17, $Gen^R$ | This study |
| WT mCherry-PLEKHA3 #T25 | WT/pCE114/pSF16.17, $Gen^R$ | This study |
| WT mCherry-PLEKHA3, VPS52-GFP #T35 | EFS30/pKG55, $Gen^R$, $Hyg^R$ | This study |
| WT mCherry-PLEKHA3, VPS52-GFP #T43 | EFS30/pKG55, $Gen^R$, $Hyg^R$ | This study |
| WT mCherry-PLEKHA3 #T29 | WT/pCE114/pSF16.17, $Gen^R$ | This study |
| WT eGFP-ML1 #T5 | WT/pBH87/pII99, $Gen^R$ | This study |
| WT eGFP-ML1 #T6 | WT/pBH87/pII99, $Gen^R$ | This study |
| WT eGFP-ML1 #T15 | WT/pBH87/pII99, $Gen^R$ | This study |
| WT ML1-mCherry #T3 | WT/pBH88/pII99, $Gen^R$ | This study |
| WT ML1-mCherry #T6 | WT/pBH88/pII99, $Gen^R$ | This study |
| WT ML1-mCherry #T12 | WT/pBH88/pII99, $Gen^R$ | This study |
| E2368 PLC-GFP #T8 | E2368/pCE105/pSF16.17, $Gen^R$ | This study |
| E2368 PLC-GFP #T10 | E2368/pCE105/pSF16.17, $Gen^R$ | This study |
| E2368 mCherry-PLC #T5 | E2368/pCE110/pSF16.17, $Gen^R$ | This study |
| E2368 mCherry-PLC #T7 | E2368/pCE110/pSF16.17, $Gen^R$ | This study |
| E2368 BTK-GFP #T5 | E2368/pCE107/pSF16.17, $Gen^R$ | This study |
| E2368 mCherry-BTK #T1 | E2368/pCE112/pSF16.17, $Gen^R$ | This study |
| E2368 HGS-GFP #T7 | E2368/pCE106/pSF16.17, $Gen^R$ | This study |
| E2368 HGS-GFP #T36 | E2368/pCE106/pSF16.17, $Gen^R$ | This study |
| E2368 HGS-GFP #T44 | E2368/pCE106/pSF16.17, $Gen^R$ | This study |
| E2368 mCherry-HGS #T19 | E2368/pCE111/pSF16.17, $Gen^R$ | This study |
| E2368 mCherry-HGS #T29 | E2368/pCE111/pSF16.17, $Gen^R$ | This study |
| E2368 mCherry-HGS #T32 | E2368/pCE111/pSF16.17, $Gen^R$ | This study |
| E2368 PLEKHA2-GFP #T7 | E2368/pCE108/pSF16.17, $Gen^R$ | This study |
| E2368 PLEKHA2-GFP #T48 | E2368/pCE108/pSF16.17, $Gen^R$ | This study |
| E2368 PLEKHA2-GFP #T58 | E2368/pCE108/pSF16.17, $Gen^R$ | This study |
| E2368 mCherry-PLEKHA2 #T6 | E2368/pCE113/pSF16.17, $Gen^R$ | This study |
| E2368 mCherry-PLEKHA2 #T23 | E2368/pCE113/pSF16.17, $Gen^R$ | This study |
| E2368 mCherry-PLEKHA2 #T76 | E2368/pCE113/pSF16.17, $Gen^R$ | This study |
| E2368 PLEKHA3-GFP #T2 | E2368/pCE109/pSF16.17, $Gen^R$ | This study |
| E2368 mCherry-PLEKHA3 #T3 | E2368/pCE114/pSF16.17, $Gen^R$ | This study |
| E2368 mCherry-PLEKHA3 #T16 | E2368/pCE114/pSF16.17, $Gen^R$ | This study |

| | | |
|---|---|---|
| E2368 mCherry-PLEKHA3 #T22 | E2368/pCE114/pSF16.17, Gen$^{R}$ | This study |
| E2368 GFP #T5 | E2368/pCE125, pSF16.17, Gen$^{R}$ | This study |
| E2368 GFP #T6 | E2368/pCE125, pSF16.17, Gen$^{R}$ | This study |
| E2368 GFP #T10 | E2368/pCE125, pSF16.17, Gen$^{R}$ | This study |
| E2368 GFP #T19 | E2368/pCE125, pSF16.17, Gen$^{R}$ | This study |
| E2368 mCherry #T7 | E2368/pCE126, pSF16.17, Gen$^{R}$ | This study |
| ***F. oxysporum*** | | |
| 4287 | Wild-type *Fusarium oxysporum*, Race 2 | |
| Δ*tepA* #3 | Δ*tepA:hph;* Hyg$^{R}$ | This study |
| Δ*tepA* #14 | Δ*tepA:hph;* Hyg$^{R}$ | This study |
| Δ*tepA* #25 | Δ*tepA:hph;* Hyg$^{R}$ | This study |
| Δ*tepA*/*tepA* #2 | Δ*tepA:hph; tepA; pHleo;* Hyg$^{R}$ Phleo$^{R}$ | This study |
| **Plasmid** | **Characteristics** | **Reference** |
| PN1862 (pSF15.15) | pSP72 containing 1.4-kb *Hind*III P*trpC-hph* from pCB1004 cloned into *Sma*I site. Amp$^{R}$; Hyg$^{R}$; *Nco*I-free P*trpC-hph* | S. Foster |
| PN (pSF16.17) | P*trpC-nptII*-T*trpC*; AmpR; Gen$^{R}$ | (Saikia and Scott, 2009) |
| PN4183 (pRS426) | *ori*(f1)-l*acZ*-T7 promoter-MCS (*Kpn*I-*Sac*I)-T3 promoter-*lacI*-*ori*(pMB1)-amp$^{R}$-*ori* (2 micron), *URA3;* Amp$^{R}$ | (Christianson et al., 1992) |
| PN1687 (pII99) | P*trpC-nptII*-T*trpC*, Amp$^{R}$/Gen$^{R}$ | (Lara-Ortiz et al., 2003) |
| PN4241 (pBV579) | pAN583 containing a 0.1 kb *Bsr*GI/*Bam*HI fragment containing NLS (three tandem repeats of the nuclear localisation signal from simian virus large T-antigen) from pEBFP2-Nuc | (Khang et al., 2010) |
| pPN82 | pBlueScriptII® KS(+) containing 1.4-kb *Hind*III P*trpC-hph* and P*gpdA*-eGFP-T*trp*C; Amp$^{R}$ /Hyg$^{R}$ | (Tanaka et al., 2006) |
| pKG55 | pPN94 containing P*tef-vps52-egfp*-T*trpC* (*vps52*: 2167 bp), Hyg$^{R}$ | Green, pers. comm. |
| pCE101 | pRS426 containing a P*gdpA-mssD*-T*trpC* overexpression construct | |
| pCE105 | pRS426 containing a P*gpdA*-PH$_{PLC-}$[illegible]$_{1}$-5Gly-eGFP-T*trpC* insert | This study |
| pCE107 | pRS426 containing a P*gpdA*-PH$_{BTK-}$5Gly-eGFP-T*trpC* insert | This study |
| pCE105 | pRS426 containing P*gpdA*- PLC PH domain-eGFP-T*trpC*; Amp$^{R}$ | This study |
| pCE106 | pRS426 containing P*gpdA*- HGS FYVE domain-eGFP-T*trpC*; Amp$^{R}$ | This study |
| pCE107 | pRS426 containing P*gpdA*- BTK PH domain-eGFP-T*trpC*; Amp$^{R}$ | This study |
| pCE108 | pRS426 containing P*gpdA*- PLEKHA2 PH domain-eGFP-T*trpC*; Amp$^{R}$ | This study |
| pCE109 | pRS426 containing P*gpdA*- PLEKHA3 PH domain-eGFP-T*trpC*; A Amp$^{R}$ | This study |
| pCE110 | pRS426 containing P*gpdA*-mCherry-PLC PH domain-T*trpC*; Amp$^{R}$ | This study |
| pCE111 | pRS426 containing P*gpdA*-mCherry-HGS FYVE domain-T*trpC*; Amp$^{R}$ | This study |
| pCE112 | pRS426 containing P*gpdA*-mCherry-BTK PH domain-T*trpC*; Amp$^{R}$ | This study |
| pCE113 | pRS426 containing P*gpdA*-mCherry PLEKHA2 PH domain-T*trpC*; Amp$^{R}$ | This study |
| pCE114 | pRS426 containing P*gpdA*-mCherry- PLEKHA3 PH domain-T*trpC*; Amp$^{R}$ | This study |
| pCE122 | pRS426 containing a P*gdpA-tepA*-T*trpC* overexpression construct | This study |
| pCE25 | pRS426 containing a P*gpdA*-eGFP-T*trpC* insert | This study |
| pCE126 | pRS426 containing a P*gpdA*-mCherry-T*trpC* insert | This study |
| pBH16 | pBH12 containing mCherry-NLS amplified from pBV579 | This study |
| pBH12 | pPN94 containing 723 bp *B. cinerea tub* terminator insert amplified from pNR1 downstream of *hph*; Hyg$^{R}$ | (Hassing et al., 2019) |
| pBH28 | pBH12 containing *nptII* replacing *hph* and *egfp*; Gen$^{R}$ | (Hassing et al., 2019) |
| pBH41 | pBH28 containing P*gpdA-egfp-tepA*-T*trpC*, Gen$^{R}$ | This study |
| pBH42 | pBH12 containing P*gpdA-tepA-mCherry*-T*trpC*, Hyg$^{R}$ | This study |
| pBH43 | pRS426 containing the *tepA* deletion construct, Hyg$^{R}$ | This study |

| | | |
|---|---|---|
| pBH72 | pBH12 containing P*tef-egfp-mssD*-T*trpC*, Hyg$^R$ | This study |
| pBH87 | pRS426 containing P*gpdA*- tandem repeat of ML1(1-86 bp) lipid binding domain-eGFP-T*trpC*; Amp$^R$ | This study |
| pBH88 | pRS426 containing P*gpdA*-mCherry- tandem repeat of ML1(1-86 bp) lipid binding domain-T*trpC*; Amp$^R$ | This study |

### Table S2: Primers used in this study.

| Name | Sequence (5'-3') | Purpose |
|---|---|---|
| tep5 | ACAGCTACCCCGCTTGAGCAGACATCACCATGGCCTCGTTGTTGCGGCAG | *tepA* for overexpression construct |
| tep6 | AGATTCGTCAAGCTGTTTGATGATTTCAGTTATTTCTGGCCATTCCCAAC | *tepA* for overexpression construct |
| BH60 | GTTGACGGCAATTTCGATG | verification of deletion |
| BH75 | CAGTGAGCGAGGAAGCGGAAGGCTTGCTTAGCTTGATATCTG | pBH12 amplification |
| BH76 | CTGAAATCATCAAACAGCTTG | pBH12 amplification |
| BH79 | GGAGGTGGAGGTTCTGGTGGAGGTGGATCTATGGTGAGCAAGGGCGAGGA | *mCherry* with linker forward |
| BH114 | GTGACACTATAGAACTCGACGAATTCCCTTGTATCTCTACA | Amplification of P*gdpA+egfp* forward |
| BH191 | CATAGATCCACCTCCACCAGAACCTCCACCTCCCTTGTACAGCTCGTCCATGC | Amplification of P*gdpA+egfp* with linker reverse |
| BH193 | GTCGAGTTCTATAGTGTCACC | Amplification of vector backbone |
| BH195 | GGAGGTGGAGGTTCTGGTGGAGGTGGATCTATGGCCTCGTTGTTGCGGCAG | Forward amplification of *tepA* with linker |
| BH196 | GGTGATGTCTGCTCAAGCGG | Amplification of vector backbone |
| BH197 | CTTCCGCTTCCTCGCTCACTG | Amplification of vector backbone |
| BH199 | CAAGCTGTTTGATGATTTCAGTTAAGATCTGTACAGCTCGTCCATGCCG | *mCherry* reverse overhang to pBH12 |
| BH203 | CCGCTTGAGCAGACATCACCATGGCCTCGTTGTTGCGGCAG | *tepA* forward, overhang to P*gpdA* |
| BH204 | CATAGATCCACCTCCACCAGAACCTCCACCTCCTTTCTGGCCATTCCCAACCAAG | *tepA* reverse, overhang to *mCherry* |
| BH208 | GGGTTTTCCCAGTCACGACATCGATCTTCCGCGACGCAGAGTACAACT | *tepA* 3' flank for deletion |
| BH209 | CCTTCAATATCAGTTCCAAGCTGTAGATTTTCTCATTGTTTC | *tepA* 3' flank for deletion |
| BH210 | CGTCCGAGGGCAAAGGAATAGGCCTGCGTAGGAAAAGCGAGATTATG | *tepA* 5' flank for deletion |
| BH211 | CAATTTCACACAGGAAACAGCATCGATGTCAACAAGTCAAGACACGCA | *tepA* 5' flank for deletion |
| BH212 | TTACAGAGACCGAGGCGATC | verification of deletion |
| BH213 | CGAGAGAAATCATTGCCCGG | verification of deletion |
| BH214 | GAGAGAGAGAGAGAGAGAGAG | verification of deletion |
| BH215 | CTTTGCTCTCCCCGTTGTC | verification of deletion |
| BH285 | GTGCACCTATTTCCACAAG | verification of deletion |
| BH411 | ATGGCCACCCCAGCAGGTAG | *ML1* for |
| BH412 | GAGCATGAGCTTGCAGGGCTT | *ML1* rev |
| BH413 | AAGCCCTGCAAGCTCATGCTCGGAGGAGGAGGAGGAATGGTGAGCAAG | *egfp* for with overhang to *ML1* |
| BH414 | CTACCTGCTGGGGTGGCCATGGTGATGTCTGCTCAAGCGG | P*gpdA* rev with overhang to *ML1* |
| BH415 | AAGCCCTGCAAGCTCATGCTCTAGCTGAAATCATCAAACAGCTTG | T*trpC* for with overhang to *ML1* |
| BH416 | CACAGGAGGTACTAGACTACCTTTCATCCTACATAAATAGACG | pRS426 rev |
| BH417 | GTAGTCTAGTACCTCCTGTGATATTATCCCATTCCATG | pRS426 for |
| BH418 | CTACCTGCTGGGGTGGCCATTCCTCCTCCTCCTCCCTTGTACAGCTCGTCCATGC | *mCherry* rev with overhang to ML1 |
| TtrpC-F | CTGAAATCATCAAACAGCTTGACG | T*trpC* for overexpression construct |
| pRS426-TtrpC-R | GCGGATAACAATTTCACACAGGAAACAGCCCATCTTAGTAGGAATGATTTTCG | T*trpC* for overexpression construct |

| | | |
|---|---|---|
| pRS426-Pgdp-F | GTAACGCCAGGGTTTTCCCAGTCACGACGAATTCCCTTGTATCTCTACA | P*gpdA* for overexpression construct |
| Pgdp-R | GGTGATGTCTGCTCAAGCGGGGTAG | P*gpdA* for overexpression construct |
| pRS426_F | GCTGTTTCCTGTGTGAAATTG | pRS426 backbone |
| pRS426_R | GGGTTTTCCCAGTCACGAC | pRS426 backbone |
| hph_F | AGCTTGGAACTGATATTGAAGG | *hph* for deletion constructs |
| hph_R | CGTCCGAGGGCAAAGGAATAG | *hph* for deletion constructs |
| GlyGFP-F | GGAGGAGGAGGAGGAATGGTGAGCAAGGGCGAGGAG | *egfp* for molecular probe construct |
| mCherry-F | ACAGCTACCCCGCTTGAGCAGACATCACCATGGTGAGCAAGGGCGAGGAG | *mCherry* for molecular probe construct |
| hgs1 | AGATTCGTCAAGCTGTTTGATGATTTCAGCTACTTCTTGTTGAGCTGCTC | *hgs* for mCherry molecular probe |
| hgs2 | ACAGCTACCCCGCTTGAGCAGACATCACCATGGAGCGCGCCCCCGACTGGGTCG | *hgs* for eGFP molecular probe |
| hgs3 | CCCTTGCTCACCATTCCTCCTCCTCCTCCCTTCTTGTTGAGCTGCTCGTAGC | *hgs* for eGFP molecular probe |
| plc1 | AGATTCGTCAAGCTGTTTGATGATTTCAGCTACTGGCGCTGGTCCATGC | *plc* for mCherry molecular probe |
| plc2 | ACAGCTACCCCGCTTGAGCAGACATCACCATGCTCCAGGACGACCCCGACCTCC | *plcs* for eGFP molecular probe |
| plc3 | CCCTTGCTCACCATTCCTCCTCCTCCTCCCTGGCGCTGGTCCATGCTGC | *plc* for eGFP molecular probe |
| btk1 | AGATTCGTCAAGCTGTTTGATGATTTCAGCTACTCGAGGATCTGGCAGCC | *btk* for mCherry molecular probe |
| btk2 | ACAGCTACCCCGCTTGAGCAGACATCACCATGGCCGTCATCCTCGAGAGCATC | *btk* for eGFP molecular probe |
| btk3 | CTTGCTCACCATTCCTCCTCCTCCTCCCTCGAGGATCTGGCAGCCCATGG | *btk* for eGFP molecular probe |
| plekha2-1 | AGATTCGTCAAGCTGTTTGATGATTTCAGCTAGGGGTGGCACTTGAGG | *plekha2* for mCherry molecular probe |
| plekha2-2 | ACAGCTACCCCGCTTGAGCAGACATCACCATGACCGGCCCCCCCCTCATCAAG | *plekha2* for eGFP molecular probe |
| plekha2-3 | GCCCTTGCTCACCATTCCTCCTCCTCCTCCGGGGTGGCACTTGAGGGCCTG | *plekha2* for eGFP molecular probe |
| plekha3-1 | AGATTCGTCAAGCTGTTTGATGATTTCAGCTAGGTGAGGCAGGCCTTGC | *plekha3* for mCherry molecular probe |
| plekha3-2 | ACAGCTACCCCGCTTGAGCAGACATCACCATGGAGGGCGTCCTCTACAAG | *plekha3* for eGFP molecular probe |
| plekha3-3 | GCCCTTGCTCACCATTCCTCCTCCTCCTCCGGTGAGGCAGGCCTTGCTGC | *plekha3* for eGFP molecular probe |
| TepA-P-Fwd | ATAGTCGGTTGGTTTGTAGGG | *tepA* 5’ flank for deletion |
| TepA-P-Nested | CCCCTCGCCTTGATCCCTT | *tepA* 5’ nested for deletion |
| TepA-P-Rv | GAATGCACAGGTACACTTGTTTAGCGGTCGTCAAGTTGCGG | *tepA* reverse, overhang to *hph* |
| TepA-T-Fwd | AGGGGCTGTATTAGGTCTCGGATGTAAGACCATGAGTGTGAA | *tepA* forward, overhang to *hph* |
| TepA-T-Nested | AGTTGCTTGAGAATGTGAGGG | *tepA* 3’ nested for deletion |
| TepA-T-Rv | CCTTGTGACCTTGCGTTTTATT | *tepA* 3’ flank for deletion |
| TepA-orf-Fwd | GAGGCTGGCTTGGATTTGTG | PCR confirmation |
| gpdA15B | CGAGACCTAATACAGCCCCT | *hph* for deletion constructs |
| trpterC8B | AAACAAGTGTACCTGTGCATTC | *hph* for deletion constructs |
| gpdA16B | AGGGGCTGTATTAGGTCTCG | confirmation knockout |
| trpC4B | CCTGGGTTCGCAAAGATAATT | confirmation knockout |
| Hyg-G | CGTTGCAAGACCTGCCTGAA | *hph* for deletion constructs |
| Hyg-Y | GGATGCCTCCGCTCGAAGTA | *hph* for deletion constructs |
| **Primers used for RT-qPCR.** | | |
| BH189 | GAGGAGATTCTGGAACCGGC | *tepA* forward |
| BH190 | CCTCGGCATCATCGTCATCA | *tepA* reverse |
| AC33 | TGCGTGACAAAACTCTCCAG | *mssD* forward |

| | | |
|---|---|---|
| AC34 | AGCCTTTCTCACGTTCTCCA | *mssD* reverse |
| TC399 | AAAAAGCAACCGAATGCAAG | *EF-2* forward |
| TC400 | CGAGACGACATAACTACATGTATCAAA | *EF-2* reverse |
| TC407 | TAGCTGGCGTTATGGAAAGG | *S22* forward |
| TC408 | CGATTGTGCGACTACTACCTCA | *S22* reverse |
| YL113F | GAGAATTCCAGCCACGCTAC | *pacC* forward |
| YL113R | AACCATCACCAGGCAAAGAC | *pacC* reverse |
| YL120F | ATGCAACAAAACGTTCACGA | *hepA* forward |
| YL120R | GCTGCTTTCCTCCTCACAAG | *hepA* reverse |
| YL501R | TGAAGAGCCTCCAGGAGAAG | *LpCCR1* forward |
| YL501F | CCTCATGCTCGGATGGTAAC | *LpCCR1* reverse |
| YL502F | CTGGTACTGCTACGGGAAGG | *LpCCR1* forward |
| YL502R | TCACCACCACAAGGTCCAC | *LpCCR1* reverse |

**Methods S1: Construct design.**

For the generation of the lipid molecular probes, the mouse lipid-binding domains (LBDs) were codon-optimised for expression in fungi using codon usage information generated for *E. festucae* (Schardl et al., 2013). Codon-optimised LBDs were fused to mCherry at the N-terminus with a 6x glycine linker generated as inserts in the vector pUC57 by GenScript USA Inc. (Piscataway, USA). The LBD was then amplified from these constructs and fused to eGFP at the C-terminus with a 5x glycine linker using Gibson assembly. Gibson assembly was further used to clone the mCherry-6Gly-LBD and LBD-5Gly-eGFP fragments under the control of the *Aspergillus nidulans gpdA* promoter and *trpC* terminator, to generate either P*gpdA*-mCherry-6Gly-LBD-T*trpC* or P*gpdA*-LBD-5Gly-eGFP-T*trpC* fragments within pRS426. The ML1 lipid-binding domain was ordered as codon-optimised tandem repeat sequence from Twist Bioscience (San Francisco, USA) and amplified with BH411/BH412. For the eGFP fusion construct the backbone was amplified in two fragments from other LPD-eGFP fusion constructs with BH413/416 and BH417/414. For the mCherry fusion construct, the backbone was amplified in two fragments from other LPD-mCherry fusion constructs with BH411/418 and BH415/416. The fragments were assembled by Gibson assembly.

For the generation of the *mssD* deletion construct (pCE98), a 1208-bp PCR fragment 5' of *mssD* and a 1020-bp PCR fragment 3' of *mssD* were amplified with the mssD1/mssD2 and mssD3/mssD4 primer pairs, respectively, from Fl1 genomic DNA. For the generation of the *tepA* replacement construct (pBH43), a 1,269-bp PCR fragment 5' of *tepA* and a 1291-bp PCR fragment 3' of *tepA* were amplified from Fl1 genomic DNA with the primer pairs BH210/BH211 and BH208/BH209, respectively. The pRS426 backbone was amplified with the pRS426 backbone F/pRS426 backbone R (5505 bp) primer pair, and the hygromycin resistance cassette P*trpC-hph* (1394 bp) from pSF15.15 with the primer combination hph_F/hph_R. For the replacement of *mssD*, a split marker approach (Rahanama et al., 2017) was employed in which the *mssD* replacement fragment was amplified as two pieces (mss1/hph-split-R 2025 bp; hph-split-F/mss4 2155 bp) with a 501-bp overlap in the middle of the *hph* resistance cassette. To generate the *mssD* overexpression (OE) construct (pCE101), the 3,029-bp *mssD* coding sequence was amplified with the primer pair mss5/mss6, while for the *tepA* overexpression construct, pCE122, *tepA* (1,798 bp) was amplified from Fl1 genomic DNA with the primer combination tep5/tep6. P*gpdA* and T*trpC* were amplified with the primer pairs pRS426-Pgdp-

F/pRS426-Pgdp-R (2310 bp) and TtrpC-F/pRS426-TtrpC-R (569 bp) from pPN82 and pII99, respectively. The pRS426 backbone was amplified with the primer combination pRS426 backbone F/pRS426 backbone R (5505 bp). Subsequently, all fragments were assembled using Gibson assembly.

The C-terminal mCherry fusion protein encoding constructs were assembled from three backbone fragments and the *tepA* (pBH42) coding region. The *tepA* coding region was amplified from Fl1 genomic DNA with the primer pair BH203/BH204. The backbone fragments included: a 2,817-bp fragment amplified from pBH16 with the primer pair BH76/BH75, containing T*trpC* and a *hph* cassette, a 4,608-bp fragment amplified from pBH28 with the primer pair BH197/196 containing genes for replication in *E. coli* and P*gpd*, and a 765-bp fragment amplified from pBV579 with the primer pair BH79/BH199 encoding *mCherry*. For the *tepA* N-terminal eGFP fusion protein constructs, the backbone was amplified in two fragments from pBH28. One fragment was 4,810 bp in length and was amplified with the primer pair BH76/BH193, and the other one was 3,080 bp in length and was amplified with the primer pair pBH114/BH191. For the *mssD* N-terminal GFP fusion construct, the backbone was assembled from the same 4,810 bp fragment in addition to a 717-bp fragment (*egfp*) amplified from pBH28 with the primer pair BH263/BH191 and a 795-bp fragment (P*tef*) amplified from pBH12 with the primer pair BH254/BH77. The 1,798-bp *tepA* coding region was amplified with BH195/tep6 and the 3,028-bp *mssD* coding region with the primer pair BH259/BH260. All fusion protein constructs encode a $(GGGS)_2$ linker between the protein of interest and the fluorophore.

In *Fusarium oxysporum*, targeted gene replacement with the hygromycin resistance cassette and complementation of the mutants by co-transformation with the phleomycin resistance cassette were performed as previously reported (Lopez-Berges et al., 2010). Briefly, two fragments encompassing approximately 2 kb of 5'- and 3'-flanking regions, respectively, were amplified using PCR with primer pairs TepA-P-Fwd/TepA-P-Rv and TepA-T-Fwd/TepA-T-Rv (Supplemental Figure 10 and Supplemental Table 2). The amplified fragments were then fused to overlapping parts of the hygromycin resistance cassette previously amplified with primers gpda15B/trpC8B, using the following primer combinations: TepA-P-Nested/HygG and HygY/TepA-T-Nested. PCR reactions were routinely performed with the High Fidelity Template PCR system (Roche Diagnostics, Barcelona, Spain), using a MJ Mini personal thermal cycler (Bio-Rad, Alcobendas, Spain). The two PCR products generated were used to transform protoplasts of *F. oxysporum* wild-type strain as described. Hygromycin resistant ($Hyg^R$) transformants were subjected to two rounds of monoconidial isolation. Deletion mutants were initially identified by PCR with primer pairs TepA-P-Fwd/TrpC4B and GpdA16B/TepA-T-Rv and the homologous recombination event was confirmed by Southern analysis. For complementation of the *F. oxysporum* Δ*tepA* mutant, a 5.0-kb fragment containing the *tepA* gene including promoter and terminator regions, was amplified from wild-type genomic DNA using primer pair TepA-P-Nested/TepA-T-Nested. A 2.5-kb fragment containing the phleomycin resistance ($Phl^R$) cassette was amplified using the primers gpda15B and trpC8B. Both fragments were used in the proportion 3:1 to co-transform protoplasts of the Δ*tepA* #3 mutant strain. Phleomycin resistant transformants were subjected to two rounds of monoconidial isolation and the presence of the wild type *tepA* gene was confirmed by PCR with primer pair TepA-orf-fwd/TepA-T-Rv.
